# Native and engineered human megakaryocytic extracellular vesicles for targeted non-viral cargo delivery to blood stem cells

**DOI:** 10.1101/2023.04.11.536479

**Authors:** Samik Das, Will Thompson, E. Terry Papoutsakis

## Abstract

Native and engineered extracellular vesicles (EVs) generated from human megakaryocytes (huMkEVs) or from the human megakaryocytic cell line CHRF (CHEVs) interact with tropism delivering their cargo to both human and murine hematopoietic stem and progenitor cells (HSPCs). 24 hours after intravenous infusion of huMkMPs into NOD-*scid* IL2Rγ^null^ (NSG™) mice, they induced a nearly 50% increase in murine platelet counts relative to saline control, thus demonstrating the potential of these EVs, which can be stored frozen, for treating thrombocytopenias. PKH26-labeled huMkMPs or CHEVs localized to the HSPC-rich bone marrow preferentially interacting with murine HSPCs. Using engineered huMkEVs or CHEVs, their receptor-mediated tropism for HSPCs was explored to functionally deliver synthetic cargo, notably plasmid DNA coding for a fluorescent reporter, to murine HSPCs both *in vitro* and *in vivo.* These data demonstrate the potential of these EVs as a non-viral, HSPC-specific cargo vehicle for gene therapy applications to treat hematological diseases.

**Native and engineered human megakaryocytic extracellular vesicles for targeted non-viral cargo delivery to blood stem cells**

**(Table of Contents):**

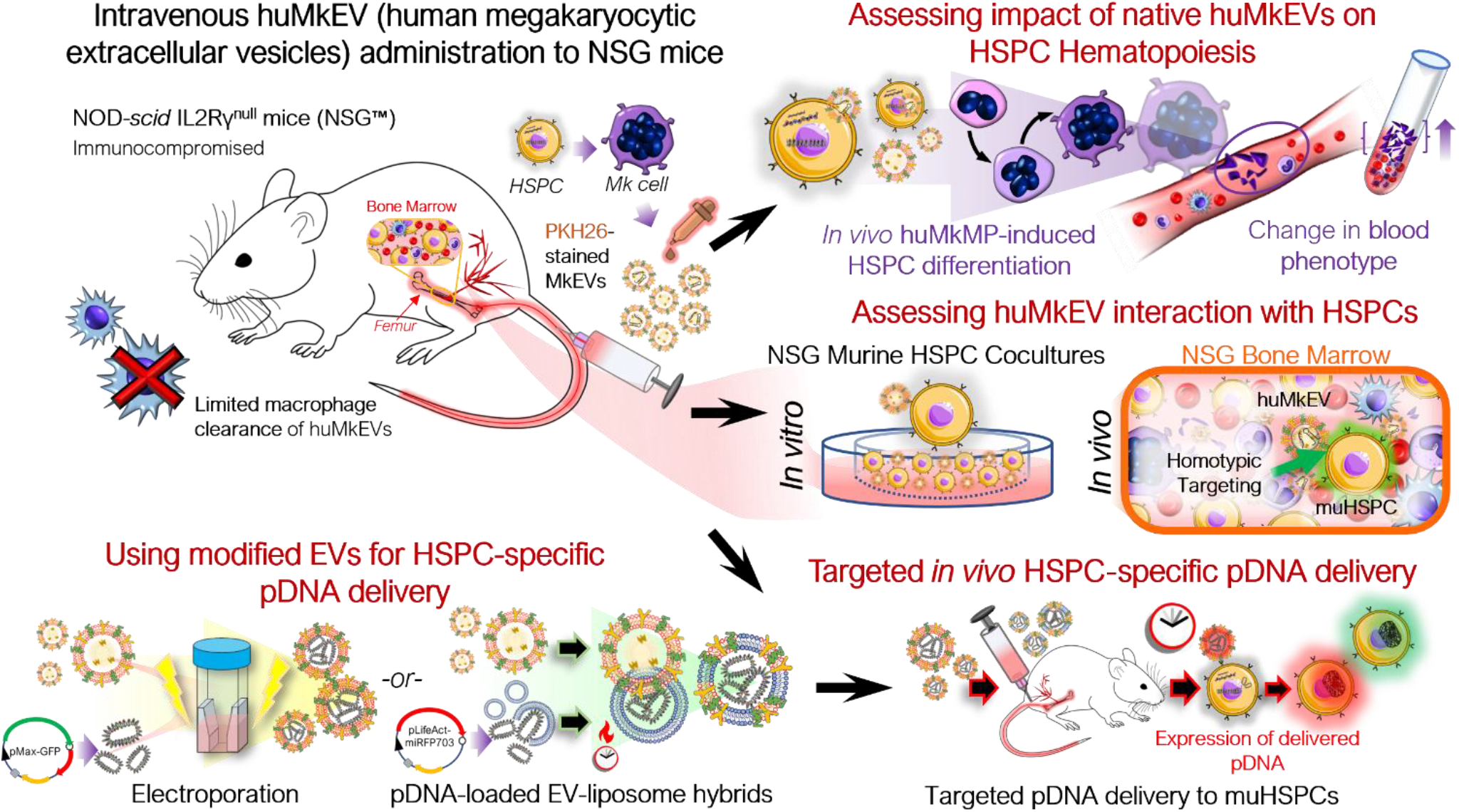

**Graphical Overview: Native and engineered human megakaryocytic extracellular vesicles (huMkEVs) for provide targeted non-viral cargo delivery to blood stem cells.**

We demonstrate that huMkEVs as a transformational cargo-delivery system to blood stem cells (hematopoietic stem and progenitor cells, HSPCs) in NOD-scid IL2Rγ^null^ (NSG™) mice. Intravenous delivery of native huMkEVs enhances *de novo* platelet biogenesis by inducing megakaryocytic differentiation of murine HSPCs, thus demonstrating the desirable strong tropism of huMkEVs for murine HSPCs. Based on this tropism, we demonstrate that engineered huMkEVs can deliver functional plasmid-DNA cargo specifically to HSPCs.

## 1. Introduction

Megakaryocytic extracellular vesicles (MkEVs), which include microparticles (MkMPs) and exosomes (MkExos), are shed from mature, platelet-producing megakaryocytes and contain endogenous RNA that play a role in the differentiation of hematopoietic stem and progenitor cells (HSPC) into megakaryocytes (**Figure 1**). ^[1, 2]^ We have previously shown that MkEVs (enriched in MkMPs, which are amongst the most abundant microparticles in circulation) interact with HSPCs with specificity/tropism and elicit megakaryopoiesis *in vitro*, without the need for the lineage specific growth factor, thrombopoietin (TPO). ^[1-3]^ Tropism for, or specificity of targeting of, HSPCs is mediated by specific huMkEV receptors. ^[3]^ We have also demonstrated that huMkEVs (human MkEVs) induce *de novo* platelet biogenesis following intravenous administration to wild type (WT; Balb/*c*) mice. ^[4]^ There was an almost 50% increase in platelet count in mice 16 hours post huMkEV administration, but a smaller increase was demonstrated 72 hours post administration due to huMkEV clearance in WT mice. ^[4]^ Despite fast clearance of labeled huMkEVs resulting in low label intensity, biodistribution studies demonstrated that administered huMkEVs preferentially localized in HSPC-rich organs and notably the bone marrow 24-hours post huMkEV administration. This suggested that some degree of huMkEV tropism to HSPCs in preserved *in vivo*. Fast clearance of huMkEVs in WT mice prevented biodistribution studies beyond 24 hours.

**Figure 1.**
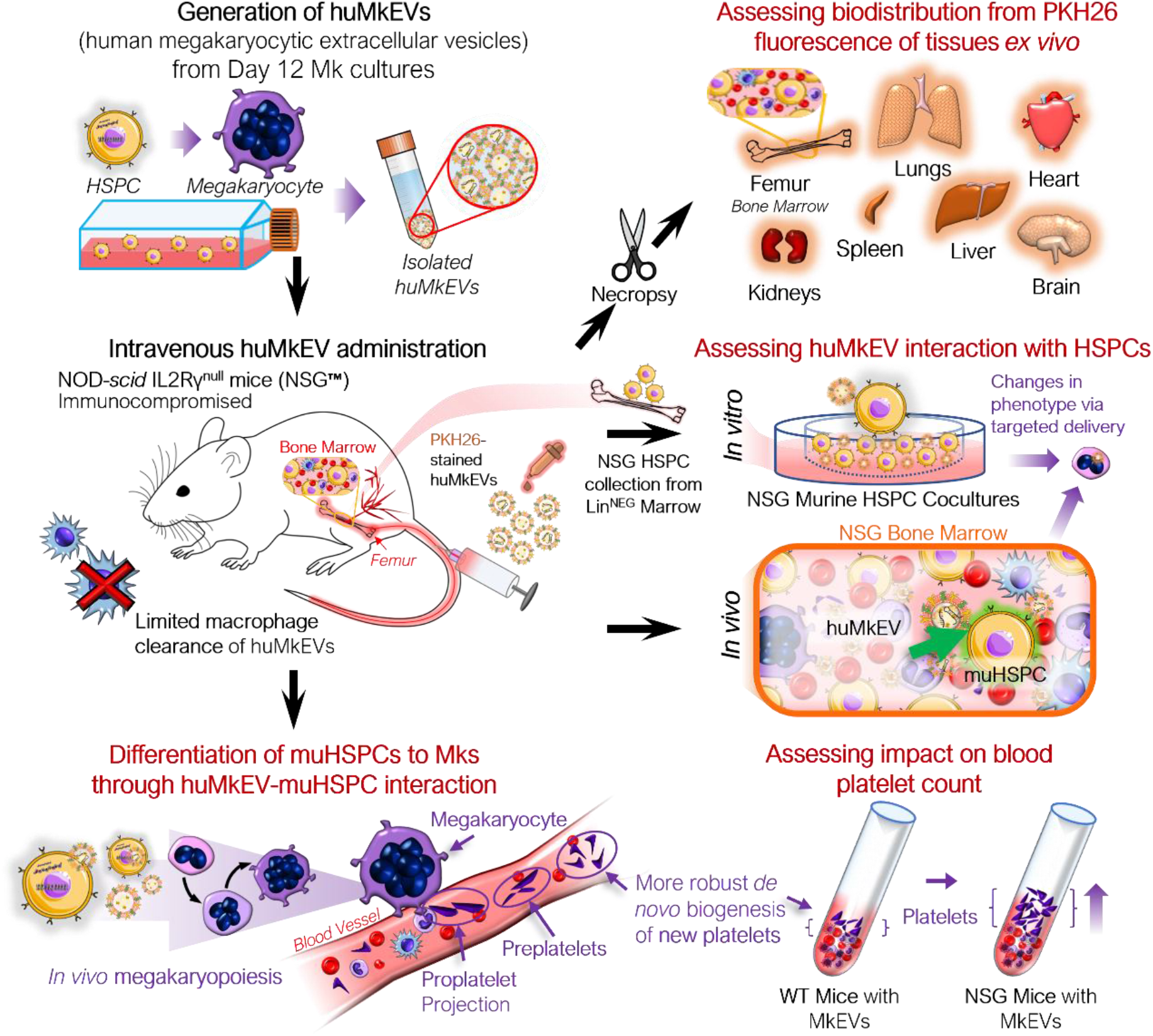
Experiment schema for assessing the *in vivo* impact of huMkEVs on NSG™ mice. The NOD-scid IL2Rγ^null^ (NSG™) mouse model was used to determine if the prior observations regarding huMkMP treatment elicit stronger responses in the NSG mice. This particular strain of mice was chosen due to its immunocompromised state devoid of functional macrophages, which could allow for prolonged exposure to huMkMPs due to extended circulation times.

Based on our studies summarized above, we aimed to develop a non-viral and non-immunogenic system for delivering cargo, including nucleic acids and proteins, to HSPCs for gene therapy and genome editing applications. Effective cargo delivery to HSPCs for gene therapy applications to treat thalassemias and other hematological abnormalities and diseases remains a major unmet need. ^[5-7]^ We have previously demonstrated that EVs contain a variety of different endogenous cargo, such as micro RNAs (miRNAs), that are dynamically exchanged between cells and are critical for determining the fate of the target cell following uptake. ^[1, 8-10]^ As an initial aim, we wanted to determine if unmodified huMkEVs could localize to and affect the phenotype of HSPCs and lay the groundwork for huMkEV-based HSPC-specific delivery of therapeutics. Thus, the ultimate goal of this study was to examine if loaded huMkEVs can functionally deliver with specificity/tropism for HSPCs, plasmid DNA, both *in vitro* and *in vivo* and demonstrate their potential for targeted therapy of genetic hematological disorders.

Given the fast clearance of huMkEVs in WT mice, the immunodeficient NOD-*scid* IL2Rγ^null^ (NSG™) mouse model offered the best option for our studies. ^[11-13]^ As NSG mice are largely devoid of functional macrophages, we hypothesized that the administered huMkEVs would reside longer in circulation and/or in murine tissues compared to immunocompetent WT mice. ^[11, 14, 15]^ If the administered huMkEVs do remain in the murine system for an extended period of time, we hypothesized that the response to infused huMkEVs will be magnified in comparison to WT mice.

As NSG mice exhibit aberrant hematopoiesis such as lower frequencies of myeloid lineage cells compared to other humanized mice, a first goal is to test the hypothesis that huMkEVs interact with murine NSG HSPCs to induce *in vitro* megakaryopoiesis and thrombopoiesis as with WT murine HSPCs. ^[16]^ This would be demonstrated by higher numbers of platelets (PLTs), pre and proplatelets (PPTs), megakaryocytes (Mks), and MkEVs. Following positive outcomes from the *in vitro* studies of this first goal, our second goal was to examine if huMkEVs also induce *in vivo* megakaryopoiesis and thrombopoiesis and if they demonstrate tropism in interacting with murine NSG HSPCs. The success of these *in vivo* studies suggested successful *in vivo* delivery of the native huMkEV cargo to murine HSPCs. Based on that, a third goal was to examine if huMkEVs loaded via electroporation with plasmid DNA coding for a fluorescent report could deliver the plasmid to murine HSPCs, both *in vitro* and *in vivo*, the latter with tropism, to demonstrate fluorescent protein expression. ^[17]^ Based on promising results from the studies of the third goal with EVs loaded via electroporation, a fourth goal was to explore a more effective method for loading huMkEVs with plasmid DNA other than using electroporation. ^[17]^ This is because electroporation results in low yields of loaded huMkEVs and may also affect the levels of the receptors that enable target specificity/tropism towards, and cargo delivery to, HSPCs. For this fourth goal, we demonstrate that the recently reported method for loading pDNA via the formation of hybrid liposome-huMkEV nanovesicles leads to successful functional delivery of pDNA to murine NSG HPSCs, both *in vitro* and *in vivo*. ^[3, 18]^

## 2. Results

### 2.1. *In vitro* studies: huMkEVs effectively interact to deliver their native cargo to NSG murine HSPCs as demonstrated by their ability to induce de novo megakaryopoiesis and platelet biogenesis

As interaction with and huMkEV cargo delivery to HSPCs is receptor mediated ^[3]^, and because NSG mice have aberrant hematopoiesis (notably in erythroid and granulocytic lineages ^[19-21]^), it could not be predicted *a priori* that their HSPCs (mu^NSG^ HSPCs) will interact with huMkEVs. Thus, prior to *in vivo* studies, it was necessary to test that huMkEVs interact and deliver their cargo to mu^NSG^ HSPCs by culturing huMkEVs with mu^NSG^ HSPCs (**Figure 1**). To collect mu^NSG^ HSPCs, femurs were collected from untreated female NSG mice (*n=3*), and bone marrow cells were isolated and subsequently lineage-depleted via MACS (Miltenyi Biotec) to collect uncommitted mu^NSG^ HSPCs and cultured as described in the methods. At Day 5, each culture was immunostained for murine CD41a, CD45, and CD117, and differentiation into megakaryocytic lineage was further tested via immunostaining cultured cells for presence of β1-tubulin (TUBB1) and von Willebrand factor (VWF), both of which are characteristic of mature platelets and megakaryocytes. ^[2, 22, 23]^

Using flow cytometry, both rhTPO and huMkEV treatment resulted in significant Increase in the fraction of CD41a+ cells (**Figure 2A**) over the untreated control. Furthermore, huMkEV treatment yielded higher ploidy CD41a^+^ cells (**Figure 2B**), indicating the development of more mature megakaryocytes in culture. ^[24]^ Both rhTPO-treated and huMkEV-treated cultures generated significantly higher counts of murine megakaryocytes, but less than 20% of the megakaryocytes from rhTPO-treated cultures exhibited ≥4N ploidies as opposed to nearly 40% of megakaryocytes from huMkEV-treated cultures exhibiting high (≥4N) ploidies (**Figure 2C**). This was further confirmed after measuring for platelet counts in each culture using CD41a^+^ staining and FSC-SSC gates optimized previously for WT mice blood platelet analysis; we observed a nearly 40-fold higher platelet count and 4-fold higher pre/proplatelet count with the huMkEV-treated cultures over each other condition (**Figure 2D**). Finally, using confocal microscopy, we observed that huMkEV-treated HSPCs visibly expressed both von Willebrand factor (vWF) and β1-tubulin (TUBB1) following 5 days of treatment, which assist with platelet coagulation and are found almost exclusively on platelets and mature megakaryocytes (**Figure 2E**). These data support the hypothesis that, *in vitro*, huMkEVs effectively interact and deliver their native cargo to mu^NSG^ HSPCs.

**Figure 2.**
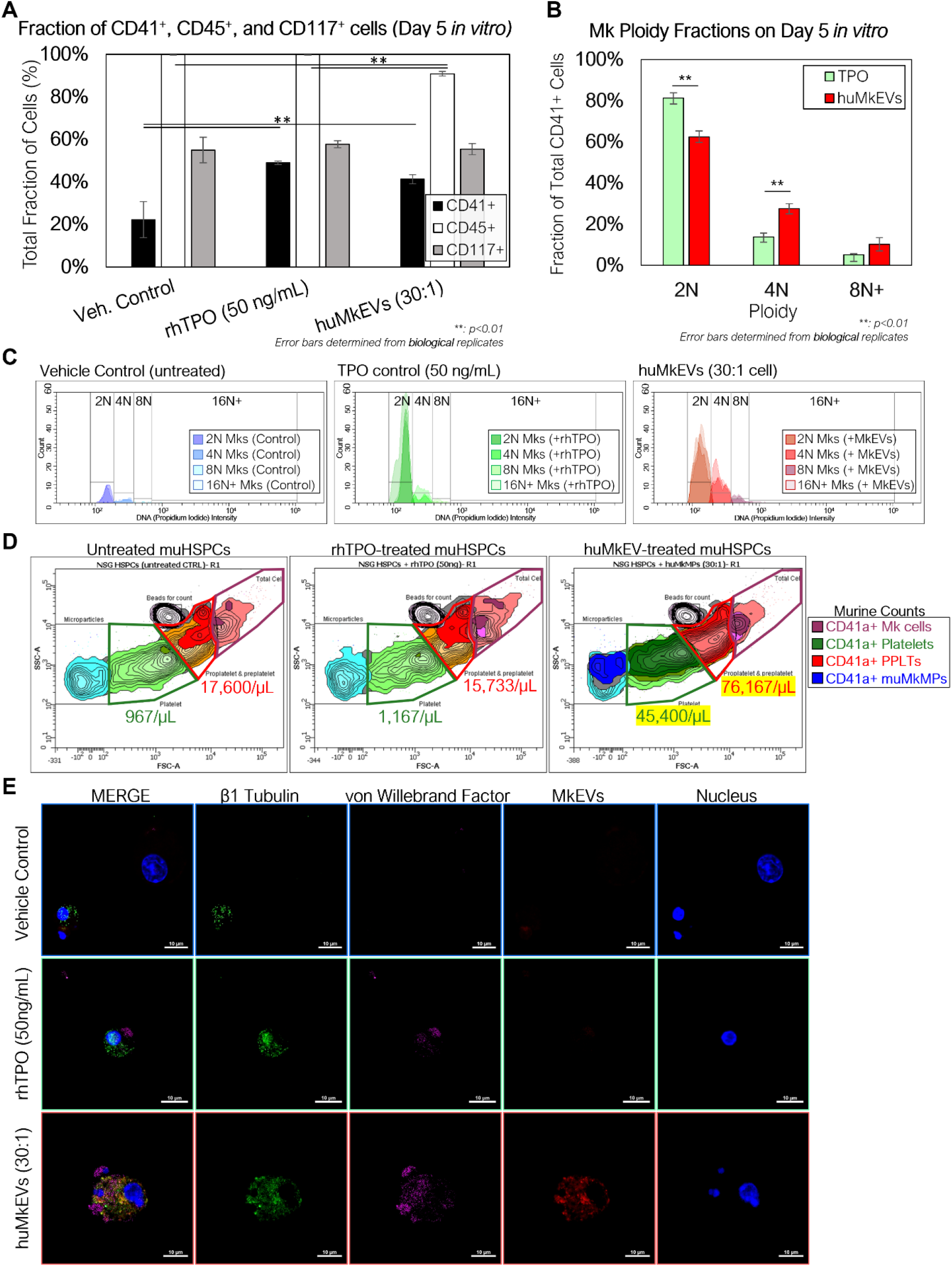
Coculture of NSG^TM^ murine HSPCs with huMkEVs in vitro induces more effective megakaryopoiesis than treatment with thrombopoietin (TPO) alone. Murine HSPCs were extracted and pooled from a set of 3 female NSG^TM^ mice, and were cultured in media containing IMDM, BIT, and 100 ng/mL stem cell factor (SCF) under the following conditions: 1) no treatment (control) 2) treatment with 50 ng/mL of rhTPO, and 3) treatment with a 30:1 ratio of PKH26-stained huMkEVs per cell. A) After 5 days, each culture was stained for murine CD41a, CD45, and CD117 and measured via flow cytometry to assess megakaryopoietic differentiation of murine HSPCs in each condition. B) Cells from each condition were permeabilized and stained with CD41a and DAPI and measured with flow cytometry to determine megakaryocytic ploidy. C) Total count-normalized histograms depicting DNA (propidium iodide) intensity of CD41a^+^ murine Mks from control (blue-shaded; left), rhTPO-treated (green-shaded; middle), and huMkEV-treated (red-shaded; right) conditions. D) Platelets, pre- and proplatelets, and microparticles were counted using calibrated fluorescent beads (1×10^6^ beads/ mL) and forward scatter (FSC), side scatter (SSC) gates set from previous wild type murine blood counts. Lighter colors correspond to a higher density of particles, while darker colors correspond to lower densities. E) The untreated (top row), TPO-treated (middle row) and huMkEV-treated (bottom row) cultures were immunostained for β1 tubulin (green; second column) and von Willebrand factor (violet; third column) to determine the degree of differentiation of the murine HSPCs to the megakaryocytic phenotype. Scale bars: 10-µm. *\*: p<0.1, **: p<0.05,* ***: *p<0.01,* ****: *p<0.001, Student’s T-test*

### 2.2. Intravenous administration of huMkEVs to NSG mice increases murine megakaryocyte and platelet biogenesis

Next, we wanted to determine if huMkEV-induced megakaryopoiesis of muHSPCs could result in *de novo* platelet biogenesis *in vivo* across a series of timepoints (**Figure 1** and **3A**). Each mouse was administered 150 µL dose of either prediluted 6×10^6^ PKH26-stained huMkEVs (*n=26* mice) or saline (PBS) solution (*n=15* mice) (**Supplemental Table S1**). The large number of mice engaged in this study was in order to examine 4 time points (4, 24, 48 and 96 hours) as each mouse is sampled at one time point only. Each dose was administered intravenously via tail vein injection. Mice were handled and kept in an aseptic environment until euthanasia. To count platelets, blood was drawn and collected from each mouse via intraperitoneal puncture immediately prior to euthanasia. After the red blood cells were lysed, we measured the concentration of murine CD41a^+^ platelets (PLTs) and pre/proplatelets (PPLTs) via flow cytometry.

**Figure 3.**
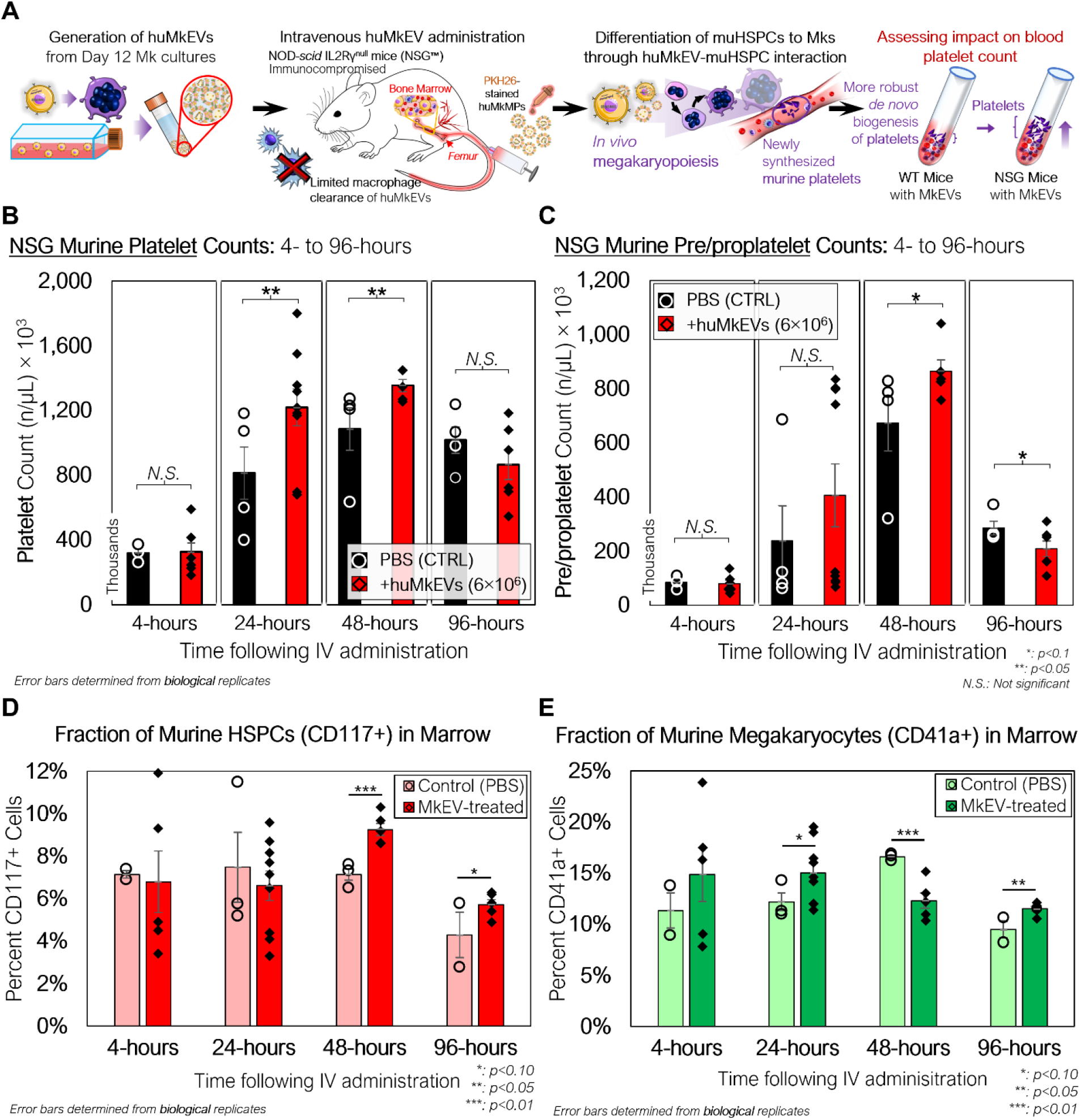
Peripheral blood platelet, pre/proplatelet, and microparticle counts and bone marrow phenotype of NSG mice treated with huMkEVs. A) Experiment schema for assessing the *in vivo* impact of huMkEVs on megakaryopoiesis in NSG™ mice. B) Platelets and C) pre/proplatelets from RBC-depleted peripheral blood were counted using calibrated fluorescent beads (1×10^6^ beads/ mL), murine CD41a staining, and forward scatter (FSC), side scatter (SSC) gates set from previous wild type murine blood counts. All values are shown as count, in thousands, per µL blood. Dashed (PBS) and solid (huMkEV) lines show average counts and trends across each timepoint. n: 3, 4, 4, 4 for PBS-treated mice and n: 6, 9, 5, 6 for huMkEV-treated mice at 4-, 24-, 48-, and 96-hours, respectively. D) Murine CD117^+^ HSPCs and E) CD41a^+^ megakaryocyte -cell fractions of flushed, RBC-depleted femoral bone marrow as determined via flow cytometry. *: p<0.10, **: p<0.05, N.S.: not significant, Student’s T-test.

There were notable differences in PLT counts between the saline-treated and huMkEV-treated mice at both 24- and 48-hours after treatment (**Figure 3B**). We observed a significant 50% boost in PLT counts in the huMkEV-treated mice after 24-hours; while still significant, the relative PLT boost diminished to 30% at 48-hours (**Figure 3B**). PPLT counts were more variable and boosts in huMkEV-induced PPLT counts were limited to 48-hours following treatment (**Figure 3C**). Differences in platelet counts subsided 96-hours following treatment, possibly indicating a return to steady-state NSG murine hematopoiesis.

As we observed a significant increase in platelet counts after 24 hours, we wanted to determine the relative fractions of CD41a^+^ (Mk) cells and Sca-1/CD117^+^ murine HSPCs within the bone marrow. We flushed out and RBC-depleted the marrow from the excised NSG femurs, and after several washes with PBS, the collected marrow was stained for CD41a^+^ and CD117^+^ cells. Relative fractions of muHSPCs in the marrow were similar at 4- and 24-hours following treatment, but muHSPC fractions at 48-hours were nearly 30% higher in the huMkEV-treated marrow than that of the saline-treated control (**Figure 3D**). CD41a^+^ cell fractions followed a different trend, with a short-term relative boost in CD41a^+^ counts measured at 24-hours in the huMkEV-treated marrow followed by a significant drop at 48-hours (**Figure 3E**). Interestingly, the relative drop in CD41a^+^ marrow cells in the huMkEV-treated mice coincides with the boost in platelet counts at 48-hours which could be a function of the maturation of murine megakaryocytes and subsequent generation of platelets. This trend was visually confirmed via confocal microscopy, with visibly greater numbers of large polyploid megakaryocytes in the marrow of huMkEV-treated mice at 24-hours (**Supplemental Figure S2**). The differences were not as notable at 48- and 96-hours (**Supplemental Figure S2**).

### 2.3. The NSG murine model enables more conclusive huMkEV biodistribution data

As the huMkEVs were pre-stained with PKH26, a strong yellow-orange lipophilic dye, PKH26 fluorescence of the excised and homogenized tissues was used to estimate huMkEV localization and biodistribution. From this assessment, the bone marrow, liver, and kidneys, exhibited immediate (4-hour) increases in mean fluorescence intensity (MFI) from the PKH26-stained huMkEVs (**Figure 4A**). When accounting for the mass of each tissue, the bone marrow exhibited the greatest increase in PKH26 MFI after 24 hours, amounting to one of the highest MFI levels per normalized tissue mass. Notably, the magnitude of the mean fluorescence intensity of PKH26 in nearly all of the tissues examined was nearly 10-20-fold higher in NSG mice in comparison to their WT counterparts ^[4]^. The NSG bone marrow exhibited comparatively high weight-normalized PKH26 fluorescence at 48- and 96-hours following treatment, but interestingly, the fluorescence from the peripheral blood increased significantly with time (**Figure 4A**).

**Figure 4.**
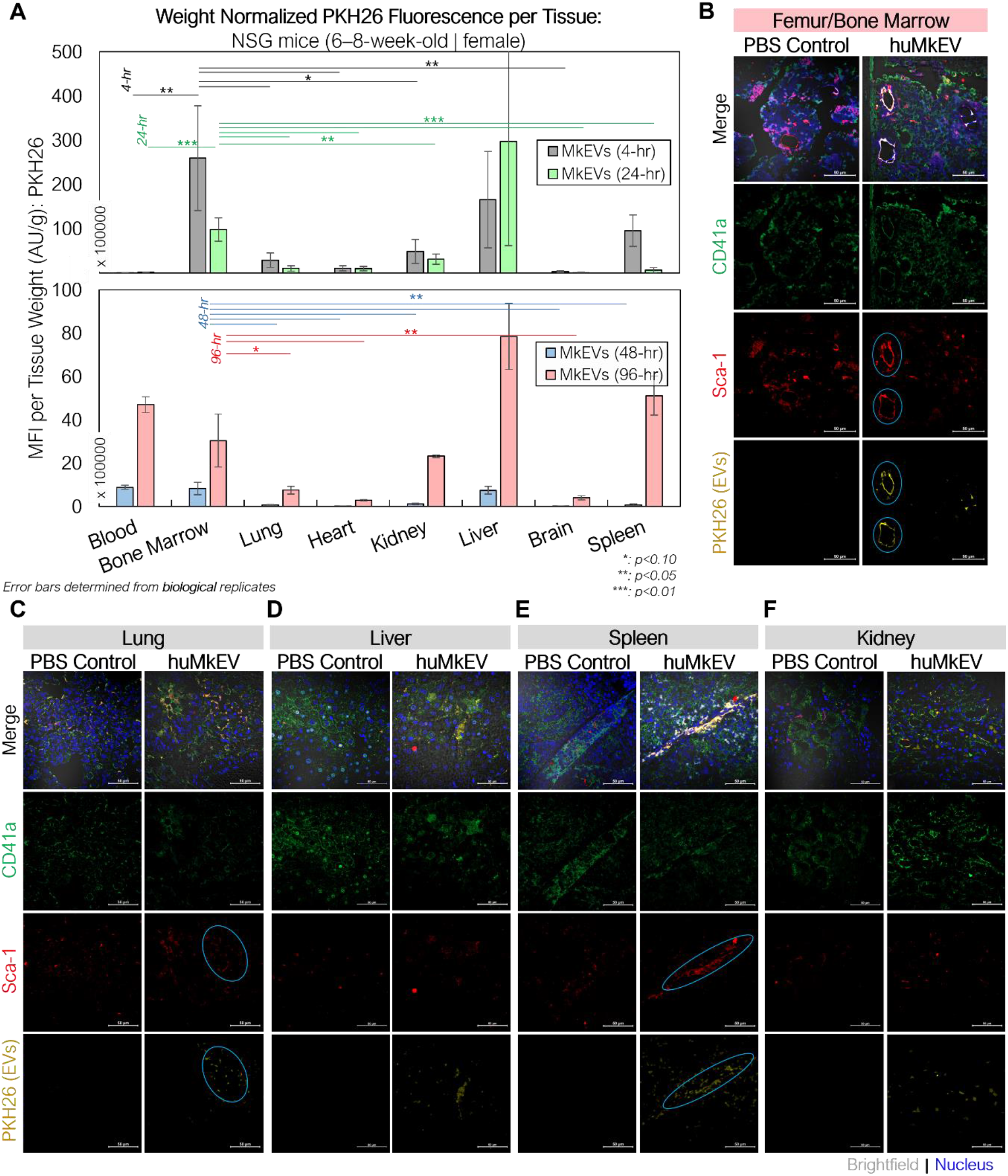
huMkEV biodistribution (MFI: AU/g) in various excised tissues and immunofluorescence assessment of bone marrow following huMkEV intravenous administration to NSG mice. **A)** Normalized mean fluorescence intensity (MFI) from each homogenized tissue reveals that huMkEVs localize to kidney, RBC^+^ bone marrow 4- and 24-hours following administration. Overall, with increasing time post administration of huMkEVs, PKH26 MFI decreases in most tissues except for peripheral blood, which shows increased MFI with increasing time. 24-hours post treatments, murine tissues from **B)** femurs, **C)** lungs, **D)** livers, **E)** spleens, and **F)** kidneys were fixed (10% neutral-buffered formalin), sectioned, immunostained and assessed for structure (gray-DIC) and presence of CD41^+^ cells (green), Sca-1^+^ cells (blue), and PKH26 fluorescence arising from administered huMkEVs. Colocalization of PKH26 fluorescence with either CD41^+^ or Sca-1^+^ cells is shown in white-orange in the merged image and circles in teal in their respective channels. Scale bars: 50-µm. *\*: p<0.1, **: p<0.05,* ***: *p<0.01, Student’s T-test*

Immunostaining the various tissues with murine CD41a and Sca-1 also helped us determine if the huMkEVs preferentially localized to these specific cell types (CD41a^+^ or Sca-1^+^ cells) within the tissue. As we have previously reported, huMkEVs targets and induces the megakaryocytic differentiation of not only the most primitive hematopoietic stem cells but also early progenitor cells. ^[2]^ Sca-1 is more abundantly expressed on both hematopoietic stem and progenitor cells than the more primitive CD117 marker. ^[25-27]^. Thus, using Sca-1, we can visualize a broader set of colocalization events with physiologically important HSPCs in tissues. After 24-hours, we observed high degrees of colocalization between the PKH26-fluorescent huMkEVs and the Sca-1^+^ murine HSPCs lining the femoral bone marrow (**Figure 4B**). The frequency of murine HSPCs was lower in the other screened tissues, such as the lungs (**Figure 4C**), liver (**Figure 4D**), spleen (**Figure 4E**), and kidneys (**Figure 4F**), but there was a strong degree of colocalization between huMkEVs and murine HSPCs in both lungs and spleen. Both huMkEVs and Sca-1 cells were at their greatest density in the periphery of the blood vessels within the vasculature of the bone marrow, which is a one of the most HSPC-rich regions of bone marrow. ^[28]^ These data demonstrate that, *in vivo,* huMkEVs retain their tropism for murine HSPCs, thus demonstrating the potential of using huMkEVs as vectors for targeted delivery of genetic cargo to HSPCs.

### 2.4. *In vitro* and *in vivo* functional plasmid DNA delivery to murine HSPCs using engineered huMkEVs

We have previously shown that plasmid DNA (pDNA) can be successfully loaded into huMkEVs via electroporation, and the pDNA-loaded huMkEVs can functionally delivered to human HSPCs (CD34^+^ cells) *in vitro* ^[17]^. Functional delivery was demonstrated by showing delivery of the pDNA to the huHSPC nucleus, and by low level expression of the plasmid encoded green fluorescence protein (GFP). Delivery to WT or NSG murine HSPCs either *in vitro* or *in vivo* has not been examined.

To test first *in vitro* delivery, we cultured lineage-depleted murine HSPCs collected from untreated NSG mice femurs with PKH26-stained huMkEVs loaded via electroporation with Cy5-labeled pDNA expressing for GFP (pMax-GFP) (**Figure 5A**). Within 24 hours of incubation, nearly 40% of the NSG HSPCs expressed GFP, indicating both successful delivery and functional expression of the pDNA. This expression sustained for 48 hours following incubation but subsided after 72 hours (**Figure 5B**).

**Figure 5.**
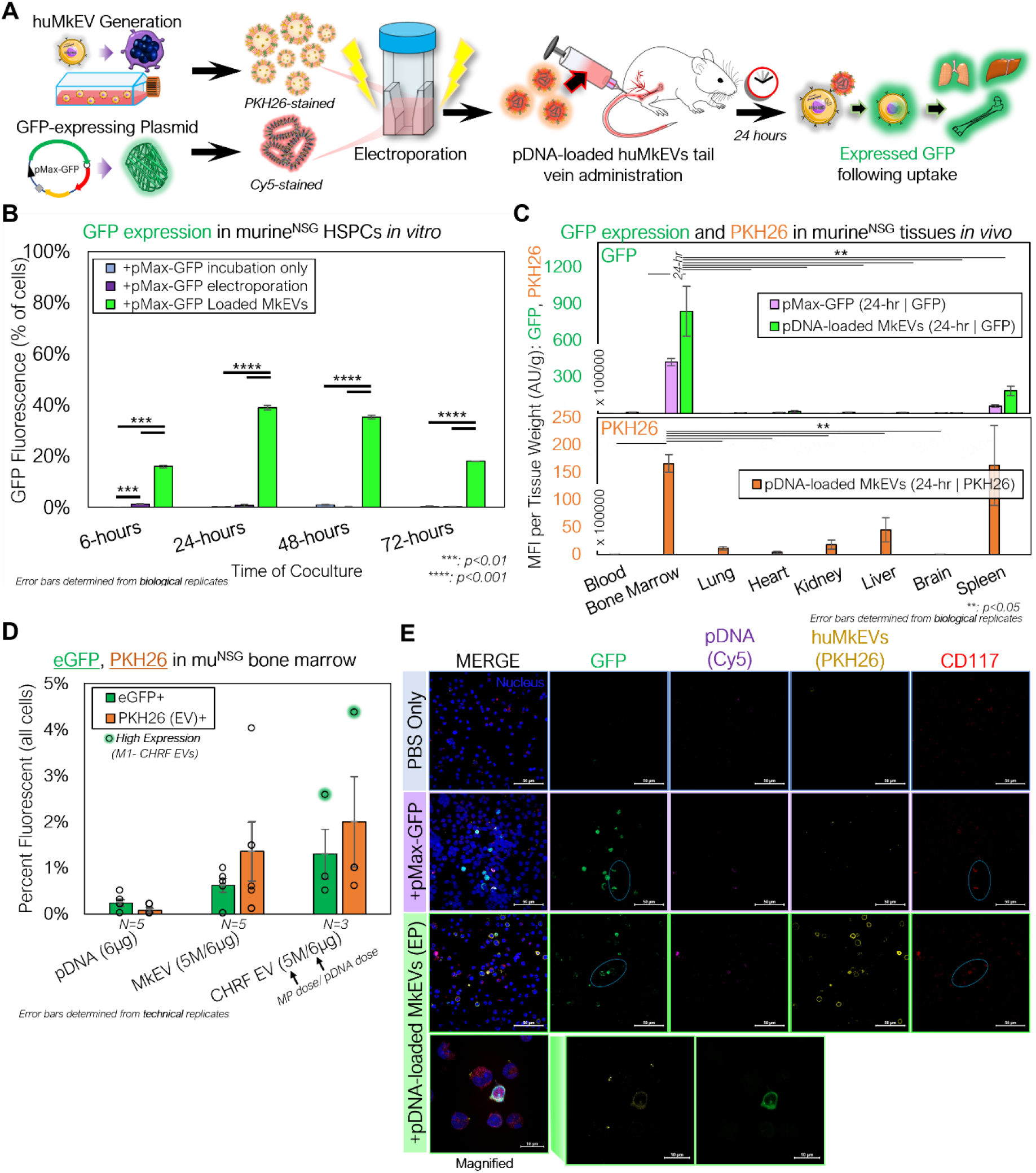
huMkEVs and CHRF EVs (CHEVs) can be electroporated with plasmid DNA for HSPC-specific nucleic acid delivery. **A)** Experimental schema for preparing pMax-GFP-loaded huMkEVs via electroporation for targeted pDNA delivery to murine HSPCs *in vitro* and *in* vivo. **B)** HSPCs from NSG mice were cultured with either ‘free’ pDNA (pMax-GFP), directly electroporated with pDNA, or cocultured with huMkEVs loaded via electroporation with pDNA, and assessed for GFP expression. **C)** 6×10^6^ pDNA-loaded huMkEVs via electroporation or an equivalent amount of ‘free’ pDNA were intravenously administered to NSG mice (*n=2*, each condition), and tissues were excised, homogenized, and assessed for GFP (top) and PKH26 fluorescence (bottom; huMkEV-treated only) after 24-hours of treatment. **D)** Flushed marrow cells from mice treated either with ‘free’ pDNA, pDNA-loaded huMkEVs, or pDNA-loaded CHEVs were assessed for GFP and PKH26 fluorescence using flow cytometry; background fluorescence was accounted for from data of untreated (PBS) mice. **E)** Flushed RBC-depleted marrow cells from the two pDNA conditions and a PBS-treated control were immunostained for CD117^+^ HSPCs (red) and screened for GFP (green), Cy5-stained pDNA (purple), and huMkMPs (yellow). Colocalization of murine HSPCs and GFP indicated with cyan circles and nuclei are shown in blue. Scale bars: 50-µm. Magnified inset shows distinct PKH26 and GFP puncta within flushed marrow cells from pDNA-loaded huMkEV-treated mice. Scale bars: 10-µm. *\*:p<0.1,**: p<0.05,* ***: *p<0.01,* ****: *p<0.001, Student’s T-test*

For *in vivo* studies, we intravenously administered prediluted 6×10^6^ PKH26-stained huMkEVs loaded with Cy5-dyed pMax-GFP via electroporation (*n= 2* mice). As a control, two mice were also treated with a dose of ‘free’ pDNA, which was equivalent to the total amount of pDNA electroporated into the huMkEVs. After 24-hours, murine tissues (bone marrow/femurs, lungs, hearts, kidneys, liver, brain, spleen, and blood) were excised, homogenized, and analyzed for GFP fluorescence. We observed that only the flushed femoral bone marrow and spleen contained GFP fluorescence following either pDNA alone or pDNA-loaded huMkEV treatment, with the marrow containing significantly higher GFP fluorescence than any other analyzed tissue (**Figure 5C**). PKH26 fluorescence, which is also a measure of localization and biodistribution of the administered huMkEVs, followed a similar biodistribution pattern (**Figure 5C**).

Similar effectiveness was demonstrated when we analyzed bone marrow cells collected from NSG mice treated with an equivalent amount of pDNA-loaded CHEVs, EVs generated from cultured megakaryoblastic CHRF-288 cells which (**Figure 5D**). Upon further analysis of the flushed bone marrow cells, we observed that the huMkEV- or CHEV-delivered pDNA resulted in greater GFP expression than pDNA alone (**Figure 5D**). Additionally, after immunostaining the flushed bone marrow cells for CD117, confocal microscopy revealed colocalization between murine HSPCs and GFP (**Figure 5E**) in mice treated with either pDNA alone or pDNA-loaded huMkEVs.

While these results demonstrated the potential of huMkEVs as a vehicle for HSPC-specific nucleic acid delivery, the low yield of electroporation-loaded EVs due to EV agglomeration prompted us to explore other methods for loading cargo into huMkEVs. Lin *et al.* developed a method of producing hybrid pDNA-loaded liposome HEK293 EV hybrids for delivering pDNA to hard-to-transfect mesenchymal stem cells. ^[18, 29]^ For these experiments, we selected the pLifeAct-miRFP703 plasmid which expresses the far-red miRFP703 which is less impacted by tissue autofluorescence. ^[30-32]^ We thus constructed miRFP703-expressing pDNA -loaded liposome-CHRF EV hybrids (CHEV hybrids; **Figure 6A**). Nanoparticle tracking analysis (NTA) of the loaded hybrid particles indicated a ∼57-nm increase in the mean diameter over the unmodified CHRF EVs (**Figure 6B)**. A change in morphology was also observed via transmission electron microscopy (TEM; **Figure 6C**). Additionally, the zeta potential of the CHEV hybrids fell between that of the CHEVs and the liposomes, further indicating hybridization between the two types of vesicles (**Figure 6D**). Greater than 90% of the pDNA was loaded into both the liposomes and the CH EV-liposome hybrids, indicating efficient encapsulation of the pDNA payload **Figure 6E**).

**Figure 6.**
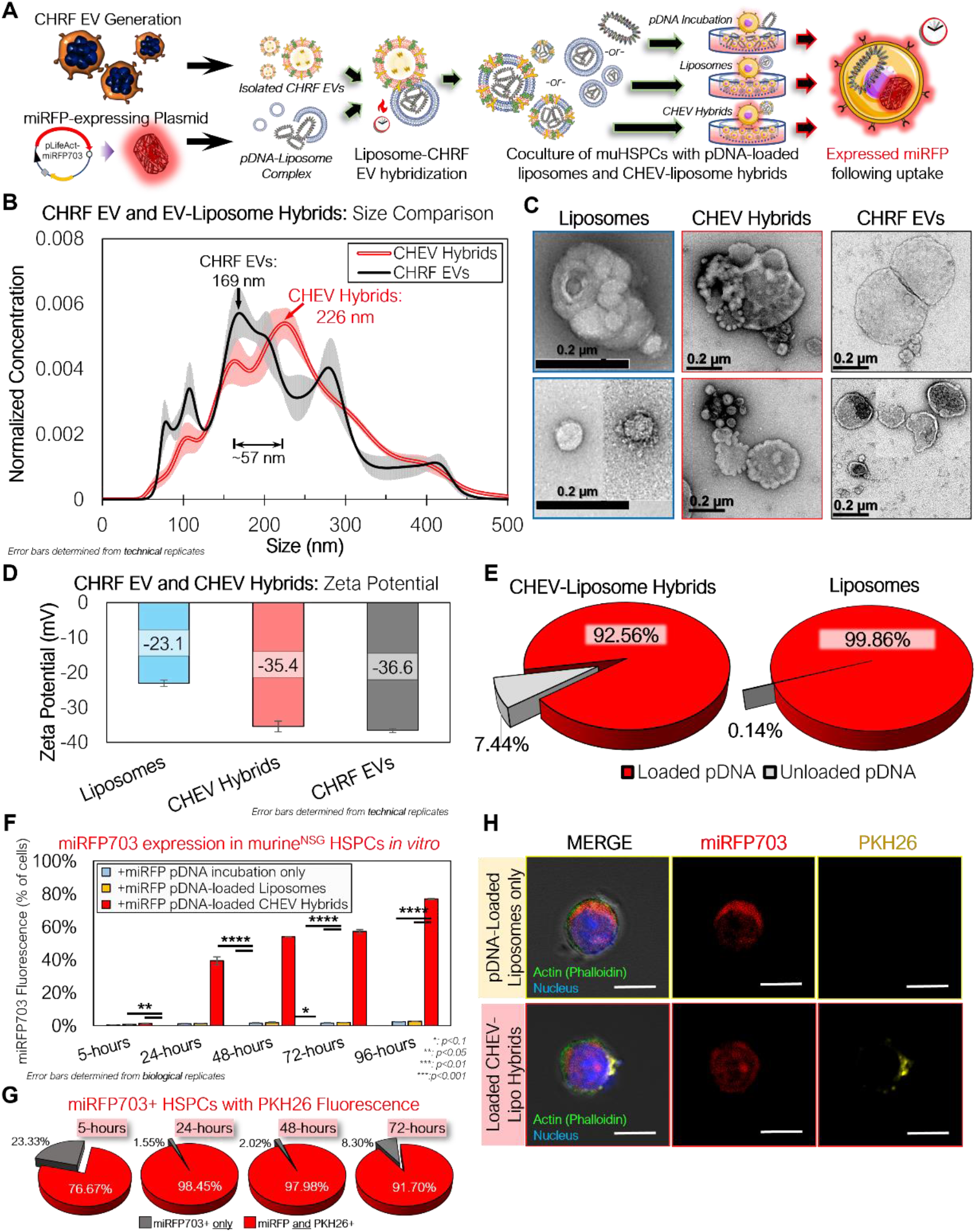
CHRF EVs (CHEVs) can be loaded with plasmid DNA via hybridization with pDNA – loaded liposomes and can be used for efficient *in vitro* nucleic acid delivery to murine HSPC. **A)** Experimental schema for preparing pLifeAct-miRFP703-loaded liposomes or CHEV-liposome hybrids for targeted pDNA delivery to murine HSPCs *in vitro.* **B)** Nanoparticle tracking analysis (NTA) of isolated CHEVs and CHEV-liposome hybrids showing an increase in diameter following hybridization. **C)** TEM images of loaded liposomes, CHEV-liposome hybrids (CHEV hybrids), and CHEVs. Scale bars: 0.2-µm/ 200-nm. **D)** Zeta potential of liposomes, CHEV hybrids, and CHEVs. **E)** Fraction of pDNA retained within the liposome or hybrid particles. **F)** Isolated NSG HSPCs were incubated with either ‘free’ (unloaded) pDNA expressing far-red miRFP703, pDNA-loaded liposomes, or pDNA-loaded CHEV hybrids and assessed for miRFP703 expression. **G)** Fraction of miRFP703^+^ HSPCs also exhibiting PKH26 fluorescence, indicating pDNA delivery via CHEV hybrids. **F)** Actin-stained (green) HSPCs from 96-hour cultures were screened for miRFP703 (red) and PKH26 fluorescence from pre-stained CHEVs to assess the presence of CHEV hybrids (yellow). *\*: p<0.1, **: p<0.05,* ***: *p<0.01,* ****: *p<0.001, N.S.: not significant, Student’s T-test*

After 24-hours of incubating CHEV hybrids with NSG HSPCs, nearly 40% of the cocultured HSPCs expressed miRFP703 (**Figure 6F**). This fraction was similar to the earlier *in vitro* coculture with the electroporated EVs at 24-hours (**Figure 5B**), but the fraction of fluorescent cells continued to climb at 48- and 72-hours with the CHEV hybrid-treated muHSPCs. Greater than 90% of the miRFP703^+^ HSPCs at 24-, 48-, and 72-hours were also PKH26^+^, indicating that the expressed pDNA was delivered via the CHEV hybrid system (**Figure 6G**). This expression was further confirmed via confocal microscopy (**Figure 6H**), with both liposome- and CHEV hybrid-treated muHSPCs exhibiting robust miRFP703 expression within the cell cytoplasm.

After confirming successful delivery of and expression from the miRFP703 plasmid *in vitro*, we used pDNA-loaded CHEV hybrids to assess targeted pDNA delivery *in vivo* (**Figure 7A).** As a control, unhybridized pDNA-loaded liposomes were prepared using the same amount of pDNA (10 µg). An equivalent volume of IMDM was prepared as an additional control. 72-hours following treatment, murine tissues were excised and analyzed for PKH26 and miRFP703 fluorescence.

**Figure 7.**
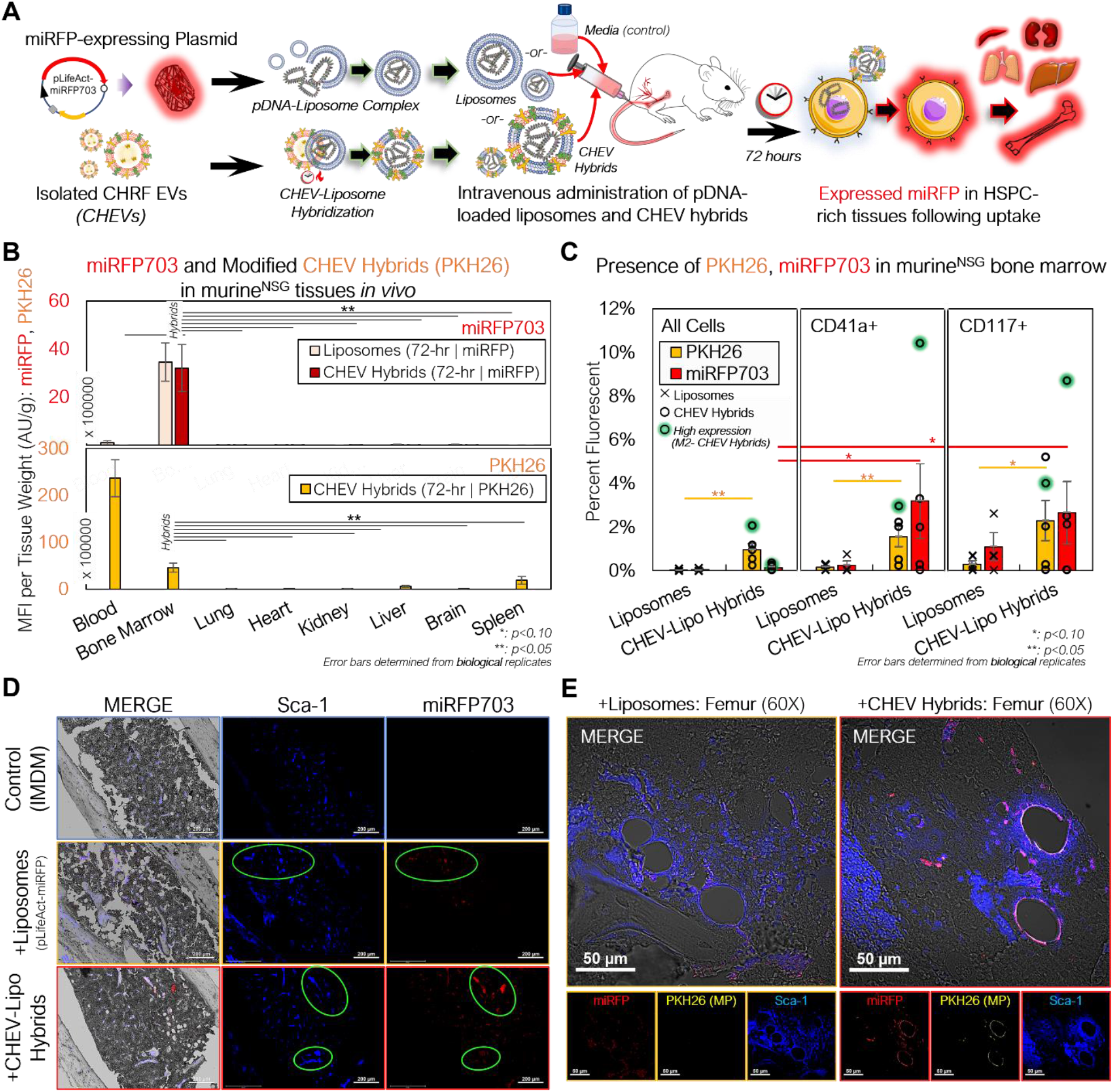
pDNA-loaded CHEV-liposome hybrids can be used for efficient HSPC-specific nucleic acid delivery *in vivo*. **a)** Experimental schema for preparing pLifeAct-miRFP703-loaded liposomes or CHEV-liposome hybrids for targeted pDNA delivery to murine HSPCs *in* vivo. **b)** NSG mice were intravenously administered either 10 µg of miRFP703-expressing pDNA complexed with liposomes (*n=3*) or pDNA-loaded liposomes further hybridized with ∼50×10^6^ PKH26-labeled CHEVs (CHEV hybrids; *n=5*), and various tissues were excised, homogenized, and assessed for PKH26 (gold) and miRFP703 (red; top) fluorescence and PKH26 biodistribution (CHEVs-only; bottom) 72-hours post particle intravenous delivery. **c)** Flushed RBC-depleted marrow cells from mice treated with either pDNA-loaded liposomes (pLifeAct-miRFP703) or CHEV hybrids and a media (IMDM)-treated control were immunostained for CD117^+^ murine HSPCs and CD41a^+^ murine Mk cells and screened for miRFP703 expression and CHEV hybrid localization (PKH26-fluorescent) via flow cytometry. High miRFP703-expressing mouse indicated in green. **d)** Femurs from 72-hour-treated mice were fixed (10% neutral-buffered formalin), sectioned, immunostained and assessed for structure (gray-DIC) and presence of Sca-1^+^ cells (blue), miRFP703 expression (red) and PKH26 arising from administered CHEV-liposome hybrids. Colocalization of murine HSPCs and miRFP703 indicated with light green circles. Scale bars: 200-µm. **e)** Magnified image of sectioned femurs from liposome- and CHEV-liposome hybrid-treated mice showing extensive miRFP expression and colocalization of expression with Sca-1^+^ HSPCs. Colocalization of Sca-1^+^ and miRFP703 shown as lavender in the merged image. Scale bars: 50-µm. *\*:p<0.1:**: p<0.05,* ***: *p<0.01,* ****: *p<0.001, Student’s T-test*

In assessing the biodistribution of the CHEV hybrids, the tissue mass normalized PKH26 fluorescence revealed a different pattern compared to our previous biodistribution studies using either unloaded huMkEVs or electroporated pDNA-loaded huMkEV-treated mice (**Figures 4A, 5C**), as fluorescence was almost exclusively limited to the peripheral blood and bone marrow (**Figure 7B**). The magnitude of PKH26 fluorescence in the peripheral blood was also 5-fold higher than the blood from huMkEV-only treated mice at any prior assessed timepoint (**Figures 4A, 5C, 7B**). This may indicate that the CHEV hybrids may circulate in the blood for extended time periods, thus explaining their limited localization in other tissues. As the PKH26 fluorescence was largely limited to the blood and bone marrow, miRFP703 was exclusively limited to the bone marrow in both the liposome and CHEV hybrid-treated mice (**Figure 7B**). Unlike the use of huMkEVs (**Figure 3**), there were no significant changes in platelet and pre/proplatelet counts between the control and either liposome- or CHEV hybrid-treated mice (**Supplemental Figures S3A & S3B**). This suggests possible dilution or loss of some of the native EV cargo (which is responsible for inducting the megakaryocytic differentiation of HSPCs) in the CHRF EVs following hybridization with the liposomes and/or differences in the endogenous cargo between MkEVs and CHEVs. ^[1, 3]^

Further analysis of the flushed bone marrow cells revealed that the CHEV hybrids retained their tropism to HSPCs *in vivo*. After immunostaining the marrow cells for murine CD117 and CD41, we found that miRFP703 fluorescence was significantly enriched in murine HSPCs (CD117^+^ cells) and megakaryocytes (CD41^+^ cells) compared to other marrow cells and to the liposome only control (**Figure 7C**). The liposome-only delivery is a measure of expression level without specific targeting, and one should consider the possibility that the clearance of loaded liposomes is slower than the clearance of the loaded hybrid particles, as these hybrids contain human membranes and proteins. While the colocalization fractions were low (∼3%) (**Figure 7C**), they demonstrate that targeting of muHSPCs by CHEV hybrids is retained. The flushed marrow cells also exhibited distinct PKH26 and miRFP703 fluorescent puncta under confocal microcopy, further confirming successful delivery of the pDNA to the marrow (**Supplemental Figure S4**).

To probe if there were any distinct colocalization patterns between miRFP expression and murine HSPCs, we immunostained the femoral bone marrow for Sca-1, which, as discussed previously, identifies more committed murine HSPCs. Both PKH26 and miRFP703 fluorescent puncta were extensively colocalized with the Sca-1^+^ marrow cells of the CHEV hybrid-treated mice, demonstrating that plasmid delivery to HSPCs using the loaded hybrid particles is feasible (**Figure 7D**). Upon further magnification, we also found extensive localization of strong miRFP fluorescence on the periphery of the blood vessels in the bone marrow vasculature of CHEV hybrid-treated mice, but to a lesser extent for liposome-treated mice, (**Figure 7E**). The periphery of bone-marrow blood vessels is an HSPC-rich region ^[28]^. We also found PKH26 fluorescence localized to these same vessel walls in the CHEV hybrid-treated marrow, although much less pronounced than for miRFP fluorescence (**Figure 7E**).

We also immunostained other tissues in addition to the femoral bone marrow, and we detected miRFP expression and some colocalization to other Sca-1^+^ cells in spleen and kidneys, albeit to a much lesser degree than the marrow (**Supplemental Figure S5**). Given the data of Figure 7C, we observed miRFP expression in CD41^+^ cells within the femurs but did not observe any distinct localization patterns in the other tissues like what we observed with Sca-1^+^ cells (**Supplemental Figure S6**).

Taken together, this data suggests the CHEV hybrids retain their tropism to HSPCs *in vivo*, further demonstrating their potential versatility as a tailorable vehicle for targeted cargo delivery to HSPCs.

## 3. Discussion

In these studies, we showed that huMkEVs robustly spur megakaryopoiesis of murine HSPCs *in vitro* and *in vivo* and can ultimately spur *de novo* biosynthesis of platelets. As we have previously suggested, given that they can be stored frozen, huMkEVs may offer a robust alternative to platelet transfusions for treating thrombocytopenias which affect millions of individuals worldwide. ^[4]^ Platelets for transfusion therapy are prepared from collected donor blood and is an expensive product in limited supply due to the short shelf life as platelets cannot be frozen. ^[33, 34]^ While there have been improvements in the handling and shelf life, banked platelets can only be stored for a maximum of 5 days and extended storage may reduce their overall clotting strength. ^[33, 34]^ By using the immunocompromised NSG strain, we were able to effectively probe these murine responses to huMkEVs with limited interference from the murine immune system, and thus better elucidate the phenotypic response to the huMkEV treatment. We have demonstrated that huMkEVs exhibit strong tropism for murine and human HSPCs, and intravenously administered huMkEVs home to the HSPC-rich bone marrow and promote megakaryopoiesis in NSG mice in relatively short timescales due to endogenous phenotype-determining cargo contained within the huMkEVs. ^[1, 8]^ When coupling the effectiveness of huMkEV-induced megakaryopoiesis and relative resilience to storage conditions, huMkEVs could be a valuable therapeutic for patients suffering from thrombocytopenia.

We have also shown that huMkEVs can be modified to carry exogenous cargo, thus laying the groundwork for a naturally-derived drug delivery vehicle for normally hard-to-transfect HSPCs. ^[6]^ We deployed two methods for exogenously loading huMkEVs and CHRF EVs with plasmid DNA: either by direct electroporation or using hybridization of these EVs with liposomes. While we were able to demonstrate the effectiveness of using these exogenous loading methods to create an EV-based vehicle for HSPC-specific functional cargo delivery, this concept can be further optimized for improved delivery efficiency to HSPCs and other cells. First, we selected an arbitrary dose of loaded hybrid particles to intravenously deliver to the NSG mice as an initial proof of concept. The data suggest that 10-20-fold higher dosages might provide a stronger measure of the potential of this delivery system. To improve targeting of HSPCs, EVs could be generated from cells overexpressing certain surface proteins and moieties, *e.g.,* ICAM-1 or MAC-1 in the case of HSPCs and may facilitate improved uptake into the target cell. ^[3, 35, 36]^ Other surface molecules, such as polyethylene glycol (PEG), can be used to ‘PEGylate’ the surface of liposomes and other particles and promote extended circulation of the administered PEGylated therapeutic *in vivo*. ^[10, 37-39]^ Further modifications to the payload and vesicle surface, such as cell-penetrating peptides, can also improve endosomal escape of the cargo, further improving intracellular trafficking of the delivered cargo.^[40, 41]^ Finally, the composition of the EV-liposome mix could be further altered to encapsulate a broader range of cargo, including larger plasmids, RNA, and proteins. Thus, both native and engineered huMkEVs have significant promise as a robust therapeutic for a broad spectrum of hematological diseases.

## 4. Conclusion

In conclusion, we have demonstrated that huMkEVs and MkEV-like CHEVs natively target and deliver functional cargo to HSPCs, which can ultimately translate to a modified HSPC phenotype. As HSPCs differentiate and commit to other blood cell lineages, MkEVs and CHEVs could facilitate targeted delivery of therapeutics to HSPCs to treat various hematological diseases.

## 5. Experimental Section

### Material sourcing

All materials were purchased from Thermo Fisher Scientific or Millipore-Sigma unless otherwise noted. All cytokines were ordered from PeproTech Inc.

### Preparation of huMkEVs from primary CD34 cells and CHRF EVs (CHEVs) from PMA-treated CHRF-288 cells

Human megakaryocytic extracellular vesicles (huMkEVs) were generated from Day 12 cultured megakaryocytes deriving from primary human CD34^+^ HSPCs using the protocols detailed by Panuganti *et. al.* ^[42]^ Briefly, CD34^+^ from different healthy donors (Fred Hutchinson Cancer Center) were pooled and cultured in IMDM medium containing 20% BIT9500 serum substitute (STEMCELL) supplemented with human LDL and a cytokine cocktail tailored to induced megakaryopoiesis (recombinant human thrombopoietin (TPO), stem cell factor (SCF) IL-3, IL-6, IL-9, and IL-11). On Day 7, CD61^+^ megakaryoblasts were enriched from the culture via MACS using anti-CD61 magnetic microbeads (Miltenyi Biotec), and the selected cells were re-cultured into flasks containing IMDM, BIT9500, TPO, SCF, and nicotinamide. On Day 12, MkEVs were isolated by first pelleting out any cells and debris at 2,000×g for 10 mins, and the supernatant was ultracentrifuged at 25,000×g for 30 mins at 4°C to collect the MkEVs.

Following isolation of the huMkEVs, the EV pellet was resuspended in 500 µL Diluent C and combined with a mix of 500 µL Diluent C and PKH26 dye. ^[43]^ After 5 minutes of incubation at ambient temperature, the EV suspension was quenched with 1.5 mL of 3% BSA (bovine serum albumin) solution, and the stained EVs were ultracentrifuged at 25,000×g for 30 mins at 4°C, followed by a second wash with 1X PBS. The PKH26-stained huMkEVs were resuspended in 100 µL 1X PBS and stored at −80°C prior to the *in vitro* cocultures and intravenous administration to mice.

To prepare CHRF EVs (CHEVs), CHRF-288 cells were expanded in CHRF media containing IMDM, 10% (v/v) heat-inactivated fetal bovine serum (FBS), 3.023 g/L NaHCO_3_ (sodium bicarbonate), and 1% (v/v) 100x antibiotic-antimycotic (αα), and incubated at 37°C, 20% O_2_, 5% CO_2_, and 85% relative humidity (rH). ^[44]^ To induce a megakaryocytic phenotype, CHRF cells were re-cultured in CHRF media supplemented with 1.5 ng/L of phorbol 12-myrsitate 13-6 acetate (PMA), seeded at a density of 3-4×10^5^ cells/mL, and incubated at 37°C for 3 days.^[45, 46]^ On Day 3, cells and debris were pelleted from the treated culture at 2,000×g for 10 mins, and EVs were isolated from the supernatant using the protocol used for isolating MkEVs.

### Preparation of NSG HSPCs for in vitro cocultures

To collect lineage-negative HSPCs from NSG mice, femurs from untreated NSG mice were collected and stored in tissue storage buffer comprising of RPMI 1640X buffer supplemented with 10% (v/v) heat-inactivated fetal bovine serum (FBS) and 1% (v/v) 100x antibiotic-antimycotic (αα) as described by Madaan *et. al.* ^[47]^ Next, muscles, connective tissues, and fat were cleared from the femur bones, and the cleaned bones were washed with 1X PBS containing 1% (v/v) αα and placed on a petri dish. After cleaning, the epiphyses were sliced from each end of the femur, and 3-mL of tissue storage buffer was flushed through the femur using a 20G needle equipped syringe to evacuate the bone marrow; the bone fragments were gently crushed to release more marrow. Finally, the evacuated marrow was transferred to a conical tube through 30 µm pre separation filter (Miltenyi Biotec) to collect the marrow cells.

To isolate the NSG HSPCs from the flushed bone marrow, the evacuated marrow cells were first pelleted at 300×g for 10 mins, and the marrow cells were resuspended in ACK (ammonium-chloride-potassium) buffer ^[4]^ to lyse any red blood cells (RBCs). After lysing, the RBC-depleted marrow cells were washed multiple times with PBS. Next, the pelleted marrow cells were incubated with direct lineage cell magnetic microbeads as described by Escobar *et. al*. ^[4]^ (Miltenyi Biotec), and the lineage-negative NSG HSPCs were collected after MACS depletion of the lineage-positive marrow cells. After several washes with IMDM, the isolated NSG HSPCs were prepared in co-culture media (80% IMDM, 20% BIT9500, 100 ng/mL SCF, 1% v/v αα) and incubated at 37°C (20% O_2_, 5% CO_2_, and 85% rH) until re-culturing.

### Exogenous loading of huMkEVs with plasmid DNA

3.2 µg of pMax-GFP pDNA (Lonza) was labeled with Cy5 (Mirus) and premixed with 5×10^6^ CHEVs and resuspended in up to 100 µL hypo-osmolar buffer; the pDNA-EV premix was transferred to a 2-mm electroporation cuvette and incubated at 37°C for 15 minutes. After incubation, the pDNA-EV-loaded cuvettes were electroporated at 200 V, 100 mA and immediately placed on ice. Next, the electroporated EVs were centrifuged at 1,000×g for 10 mins to remove any agglomerates, and the pDNA-loaded EVs were isolated from the supernatant following centrifugation at 25,000×g for 30 mins at 4°C. Finally, the pDNA-loaded EVs were resuspended in a small volume of filtered 1X PBS and stored at −80°C until use.

To prepare the pDNA-loaded liposome-EV hybrids, pLifeAct-miRFP703 was first loaded into liposomes (Lipofectamine 2000) using the protocol adapted from Thermo Fisher. ^[32]^ Briefly, 1 µg of pDNA and 2 µL of the Lipofectamine 2000 reagent were each diluted in 25 µL IMDM each. Next, each diluted aliquot was mixed, and the 50 µL mix was incubated at room temperature for 5 minutes. To hybridize the pDNA-loaded liposomes with PKH26-stained CHEVs, the hybridization protocol developed by Lin *et. al.* was adopted and modified. ^[18]^ After preparing the 50 µL pDNA-liposome complexes, 5×10^6^ EVs were dosed into each mix and incubated at 37°C for 12 hours. Following incubation, the CHEV-liposome hybrids were stored at 4°C until use. Separate 50 µL liposome complexes were prepared with 1 µg of pDNA and 2 µL of Lipofectamine 2000 as an additional liposome-only control condition.

To characterize the liposomes, CHEVs, and CHEV-liposome hybrids, samples were diluted in Millipore-filtered water, and 3 technical replicates of each diluted sample was measured using nanoparticle tracking analysis (NTA; Malvern NanoSight NS300). The particle size distribution was calculated using the NanoSight NS300 software. To measure the zeta potential, approximately 350 µL of each sample was loaded into an Anton Paar Omega cuvette, and three technical replicates were measured using the Litesizer 500 (Anton Paar). To quantify plasmid DNA loading efficiency, the loaded liposomes and CHRF EV-liposome hybrids were pelleted at 25,000×g for 30 mins at 4°C, and the pellets were resuspended in lysis buffer (Qiagen); the supernatants were retained to quantify ‘free’ or unloaded pDNA. Next, the lysate and supernatant were applied to pDNA binding columns, washed with PE wash buffer (Qiagen), and eluted using biology-grade water. Final pDNA concentration was determined using the Qubit DNA quantification kit (Invitrogen). TEM (transmission electron microscopy) images were prepared by the University of Delaware Bioimaging Center as described. ^[8]^

### Coculture of huMkEVs with isolated NSG HSPCs and ploidy, platelet

After collecting the NSG muHSPCs from the femurs of untreated mice, muHSPCs were cultured in media containing 80% (v/v) IMDM, 20% (v/v) BIT9500, 1% antibiotic-antimycotic, and 100 ng/mL rhSCF and incubated at 37°C with 20% O_2_, 5% CO_2_, and 85% rH for 24 hours. The following coculture conditions were used:

i) Untreated control: ∼7.5×10^4^ muHSPCs in 750 µL of coculture media only (80% IMDM, 20% BIT9500, 100 ng/mL rhSCF, 1% αα)
ii) +rhTPO: ∼7.5×10^4^ muHSPCs in 750 µL of coculture media supplemented with 100 ng/mL rhTPO
iii) +huMkEVs: ∼7.5×10^4^ muHSPCs in 750 µL of coculture media incubated with 30:1 (huMkEVs: muHSPCs), or 2.25×10^6^ PKH26-stained huMkEVs

Following 5 days of incubation at 37°C, 50 µL of each culture was incubated with FITC rat anti-mouse CD41a, APC rat anti-mice CD117, or APC rat anti-mice CD45 to measure levels of murine megakaryocytes, stem and progenitor cells, and non-erythrocyte differentiated hematopoietic cells, respectively. Fractions of CD41a+, CD45+, and CD117+ cells were gated against unstained controls. Following CD41a staining, platelets were counted via flow cytometry (BD FACSAria II) using FSC-SSC gating previously optimized for counting platelets in RBC-lysed murine peripheral blood. ^[4]^ Total platelet concentration was determined after diluting 10 µL of AccuCount beads (Spherotech), which were used as a standard.

To quantify the ploidy of megakaryocytes, the protocol of Lindsey *et. al.* was used. ^[48]^ Briefly, mature megakaryocytes were fixed with 4% paraformaldehyde in 1X PBS for 15 minutes at room temperature and subsequently permeabilized with 70% methanol for 1 hour at 4°C. Next, the permeabilized megakaryocytes were stained with FITC rat anti-mouse CD41a and the nuclei were further stained with 50 µg/mL propidium iodize to gauge ploidy. Finally, ploidy was assessed via flow cytometry, with ploidy counts determined from DNA content through the distinct peaks of propidium iodide fluorescence.

To assess the degree of megakaryopoiesis of the Day 5 cultures via confocal microscopy, each culture was immunostained for β1-tubulin (TUBB) and von Willebrand factor (VWF), which are highly abundant in platelets and critical for platelet adhesion, and thus, characteristic of mature megakaryocytes.^[23, 49]^ Briefly, 100 µL of each culture was seeded onto poly-L-lysine coated coverslips, fixed with 4% paraformaldehyde in 1X PBS, and blocked with blocking buffer (1X PBS, 10% normal donkey serum, 3% bovine serum albumin) for 1 hour at room temperature. Next, the coverslips were incubated with a 1:100 dilution of rabbit anti-VWF (Abcam) and 1:100 mouse anti-TUBB in blocking buffer at room temperature for 2 hours. After washing with PBS, each coverslip was subsequently incubated with a 1:1000 dilution Alexa Fluor 488 anti-rabbit IgG and 1:1000 Alexa Fluor 647 anti-mouse IgG_3_ κ at room temperature for an additional 2 hours. Finally, the immunostained coverslips were sealed onto glass slides with SlowFade mounting media with DAPI, sealed with nail polish, and stored at 4°C until imaging.

### Coculture of pDNA-loaded huMkEVs and CHEVs with NSG HSPCs in vitro for assessment of HSPC-specific cargo delivery

To assess the efficiency of using electroporated huMkEVs to deliver pMax-GFP pDNA, the following cocultures were set up:

i) Untreated control: ∼1.0×10^5^ muHSPCs in 750 µL of coculture media only (80% IMDM, 20% BIT9500, 100 ng/mL rhSCF, 1% αα)
ii) pDNA electroporation: ∼1.0×10^5^ muHSPCs electroporated with 1 µg pMax-GFP pDNA in 750 µL of coculture media
iii) +pDNA-loaded huMkEVs: ∼1.0×10^5^ muHSPCs cultured with 3×10^6^ pDNA-loaded huMkEVs (30:1 huMkEV: muHSPC) in 750 µL of coculture media

Following treatment, 50 µL of each sample and replicate were collected after 6-, 24-, 48-, and 72-hours of incubation and measured for GFP fluorescence via flow cytometry (BD FACSAria II) and gated against untreated controls.

To assess the efficiency of using CHEV-liposome hybrids to deliver pLifeAct-miRFP703 pDNA, the following cocultures were set up:

iv) Untreated control: ∼1.0×10^5^ muHSPCs in 750 µL of coculture media only (80% IMDM, 20% BIT9500, 100 ng/mL rhSCF, 1% αα)
v) +Liposomes: ∼1.0×10^5^ muHSPCs incubated with 50 µL containing 1 µg pLifeAct-miRFP703 liposome complexes in 750 µL of coculture media
vi) +CHRF EV-liposomes: ∼1.0×10^5^ muHSPCs cultured with 50 µL CHRF EV-liposome hybrids containing 1 µg pDNA in 750 µL of coculture media

To measure pDNA uptake and expression, 50 µL samples were collected after 6-, 24-, 48-, and 72-hours of incubation and measured for miRFP703 fluorescence using flow cytometry (Beckman CytoFLEX S). miRFP703-fluorescent populations were gated against untreated controls to exclude background fluorescence.

To prepare samples for screening via confocal microscopy, 100 µL of each 72-hour culture was seeded onto poly-L-lysine coverslips, fixed, and washed as described above. Next, to label the cell cytoskeleton, each coverslip was incubated with 500 µL filtered PBS containing 5 µL Alexa Fluor 488-conjugated phalloidin and incubated at room temperature for 1 hour. Finally, the prepared coverslips were washed with PBS, mounted onto slides with SlowFade mounting media with DAPI, and sealed with nail polish. Slides were stored at 4°C until imaging via confocal microscopy (Zeiss LSM 880).

### Administration of huMkEVs and collection of various tissues from NSG mice for biodistribution analysis

Prior to administering huMkEVs to the NSG mice, PKH26-stained huMkEVs were thawed, immunostained with FITC-conjugated mouse IgG anti-human CD41a (BD Lifesciences) and counted via flow cytometry (BD FACSAria II). Next, aliquots containing 5×10^6^ huMkEVs were each resuspended in 150 µL sterile filtered PBS and loaded into sterile 50 cc insulin syringes. After preparing the huMkEV samples, 6–8-week-old female NSG mice were briefly warmed and were administered with either 150 µL sterile PBS only or with 5×10^6^ huMkEVs in PBS via the tail vein. The mice were then grouped by treatment and housed in a sterile environment until necropsy and analysis 4-, 24-, 48-, or 96-hours following treatment.

After euthanasia using CO_2_ asphyxiation and cervical dislocation, the femurs, heart, lungs, spleen, liver, kidneys, and brain were excised and placed in 3 mL of cold tissue storage buffer (RPMI 1640X with 10% v/v FBS) ^[47]^ Next, each tissue was trimmed of any connective tissue and fat and rinsed with 1X PBS containing 1% αα (v/v). After rinsing and weighing, each tissue was placed in 2 mL bead mill homogenizer tubes containing 1 mL hypotonic lysis buffer and 2.8 mm ceramic beads (90 mg/bead; 6x beads/tube) To homogenize the tissues, each tube was loaded into a bead mill homogenizer (Fisher Bead Mill 24) and homogenized at 5.0 m/s for 20 seconds with 4 total cycles. The bone marrow was extracted from the femurs and processed without lineage depletion as described above, and the final bone marrow flushes were resuspended in 1-mL tissue storage buffer until analysis.

To measure PKH26 biodistribution, 2×200 µL of each tissue homogenate was loaded into an opaque-bottomed 96-well plate and checked for fluorescence (λ_excit_: 551 nm, λ_em_: 567 nm) using a microplate reader (SpectraMax i3x). Tissue autofluorescence was accounted for by subtracting the mean fluorescence of each tissue from PBS-only treated mice. After tabulating the net tissue mean fluorescence intensity, fluorescence values were normalized by the tissue weight of each individual tissue.

### Preparation of murine blood for platelet counts and phenotypic analysis

Prior to euthanasia and necropsy, approximately 100-200 µL of murine blood was extracted via cardiac puncture and collected in a K2-EDTA microtainer (BD Lifesciences). Next, 10 µL of the chelated blood was drawn into a capillary tube and placed in 990 µL ammonium oxalate RBC lysis buffer. After RBC lysis, 100 µL of the depleted blood was incubated with 2.5 µL FITC rat anti-mouse CD41a at 4°C for 15 minutes to immunostain and identify megakaryocytes, platelets, pre and proplatelets, and megakaryocytic microparticles via flow cytometry.

To quantify total platelet counts per unit of blood through flow cytometry (BDFACSAria II), 10 µL of AccuCount fluorescent beads (Spherotech) was added to each depleted blood sample following CD41a antibody incubation. Next, platelets and pre/proplatelets were gated on CD41a+ populations using FSC-SSC gating previously optimized for counting platelets in murine blood in wild-type mice. Total platelet concentration was calculated using the known concentration of the AccuCount beads as a standard, and final counts were adjusted for the initial dilution in RBC lysis buffer.

### Intravenous administration of pDNA-loaded CHRF EVs to NSG mice to determine in vivo cargo delivery to murine HSPCs

Prior to intravenous administration, PKH26-labeled huMkEVs and CHEVs were loaded with pMax-GFP and pLifeAct-miRFP703, respectively, as described above. Next, several aliquots of 6×10^6^ of the electroporated huMkEVs were resuspended 150 µL PBS; separate 5 µg aliquots of Cy5-labeled pMax-GFP were prepared in 150 µL PBS to serve as an additional control. After preparing the samples, 6–8-week-old female NSG mice were intravenously administered with either PBS only, pDNA, or pDNA-loaded huMkEVs via the tail vein, and all treated mice were grouped by their treatment condition and housed in a sterile environment for 24 hours.

Next, the treated mice were euthanized, and each tissue sample was homogenized and loaded into microplates as described above. Microplates were then measured for both PKH26 and GFP (λ_excit_: 475 nm, λ_em_: 509 nm) fluorescence to assess the huMkEV biodistribution and GFP expression, respectively. As before, tissue autofluorescence within both spectra was accounted for by subtracting the raw fluorescence of each tissue collected from PBS-treated mice. RBC-depleted bone marrow samples were seeded, fixed, and blocked onto poly-L-lysine coated coverslips as described above. Coverslips were subsequently immunostained with Alexa Fluor 594-conjugated rat IgG2b anti-mouse CD117 to label HSPCs and incubated for 2 hours at room temperature. Finally, the coverslips were mounted onto glass slides with SlowFade mounting media with DAPI, sealed, and stored at 4°C until imaging.

To treat NSG mice with the CHEV-liposome hybrids, the protocol described above was scaled to prepare doses comprised of 45×10^6^ PKH26-stained CHEVs and 9 µg pLifeAct-miRFP703 in 250 µL IMDM media as prepared. An equivalent amount of pDNA was complexed with liposomes as an additional control. Next, mice were intravenously administered via the tail vein with either 250 µL IMDM (negative control), pDNA-loaded liposome complexes, or pDNA-loaded CHEV-liposome hybrids. Mice were then grouped by their experimental condition and housed in a sterile environment for 72 hours until necropsy.

Following necropsy, tissues were homogenized and prepared as described previously. Next, PKH26 (λ_excit_: 551 nm, λ_em_: 567 nm) and miRFP703 (λ_excit_: 670 nm, λ_em_: 703 nm) fluorescence were measured via a microplate reader (SpectraMax i3x) were assessed to determine CHEV-hybrid biodistribution and pDNA expression, respectively; the raw autofluorescence of the IMDM-only treated mice was subtracted for each tissue in each of these respective channels. Net fluorescence values were subsequently normalized by the tissue weights. Blood platelet counts were estimated through flow cytometry (Beckman CytoFLEX S) as described in previous sections.

Femurs from the treated mice were flushed and RBC-lysed as described in Section 5.2. Next, flushed bone marrow cells were seeded onto poly-L-lysine coverslips, fixed, and initially blocked as described in Section 5.9 and Section 5.11. To label murine HSPCs, coverslips were immunostained with a 1:100 dilution (in blocking buffer) of rat IgG2b anti-mouse CD117 for 2 hours at room temperature. Next, the primary antibody-stained coverslips were secondarily stained with a 1:1000 dilution of Alexa Fluor 594-conjugated anti-rat for an additional 2 hours at room temperature. After immunostaining and washing, each coverslip was mounted on glass slides with SlowFade mounting media with DAPI, sealed, and stored at 4°C until imaging.

### Immunostaining bone marrow cells and select tissues for phenotype analysis via flow cytometry and confocal microscopy

Following necropsy, marrow cells were extracted from the femurs using the protocol outlined above without any lineage depletion. After collecting the marrow cell flushes, each flush was resuspended in an equivalent volume of tissue storage buffer, and approximately 50 µL of each suspension was retained for flow cytometry immunostaining prior to lysis. Next, to assess the fraction of murine megakaryocytes and HSPCs in the bone marrow, 50 µL aliquots of marrow cells were immunostained with 5 µL Alexa Fluor 488-conjugated rat IgG1 anti-mouse CD41 or APC-Cy7-conjugated rat IgG2b anti-mouse CD117 (BioLegend) and incubated at 4°C for 15 minutes. The immunostained marrow cells were next analyzed via flow cytometry (BD FACSAria II). Positively immunostained populations were gated against unstained controls. To determine uptake of huMkEVs by the marrow cells, cells were also analyzed for PKH26 fluorescence.

To prepare samples for confocal microscopy, RBCs were depleted from the flushed marrow cells by resuspending the bone marrow cells in ACK buffer for 5 minutes. Next, the RBC-depleted bone marrow cells were washed 2-3 times with 1X PBS at 300×g for 10 minutes. After pre-coating each coverslip with 200 µL 0.1% poly-L-lysine for at least 30 minutes and washing, 100 µL of each bone marrow sample was seeded, fixed onto coverslips, and blocked with blocking buffer using the protocol described above. Next, the fixed cells were immunostained with a 1:100 dilution of rabbit anti-CD41 and rat IgG2a anti-Sca-1 (Abcam) in blocking buffer to identify murine megakaryocytes and HSPCs, respectively. After incubating with the primary antibody mix for 2 hours at room temperature, each coverslip was stained with a secondary 1:1000 dilution of Alexa Fluor 488-conjugated anti-rabbit and Alexa Fluor 647-conjugated anti-rat for an additional 2 hours at room temperature. Finally, following several washes with PBS, each coverslip was sealed onto glass slides with SlowFade mounting media with DAPI, sealed with nail polish, and stored at 4°C until imaging.

For histological tissue analysis and immunofluorescence staining, samples collected from 4- and 24-hour huMkEV treated mice were processed at the Nemours Histology Core facility (Wilmington, DE). Samples collected from NSG mice treated with the pDNA-loaded liposomes and CHEV-liposomes were processed at the University of Delaware Histology Core facility (Newark, DE). Briefly, femurs, lungs, liver, spleen, and kidneys were fixed with 10% neutral-buffered formalin for several days. Next, after decalcification of the femur, tissues were imbedded in paraffin, sectioned, mounted on slides, and deparaffinized prior to immunostaining.

For the prepared samples collected from the huMkEV-only treated mice (and PBS control), sectioned tissues were immunostained with rabbit anti-mouse CD41 and rat IgG2b anti-mouse Sca-1 to label murine megakaryocytes and HSPCs, respectively; cell nuclei were stained with DAPI. The immunostained tissues were secondarily stained with Alexa Fluor 488 anti-rabbit and Alexa Fluor 647 anti-rat, to label CD41^+^ and Sca-1^+^ cells, respectively. Finally, the prepared samples were imaged via confocal microscopy (Zeiss LSM 880).

For the samples collected from mice treated with pDNA-loaded liposomes or CHEV-liposome hybrids, each sectioned tissue was either immunostained with rabbit anti-mouse CD41 -or- rat IgG2b anti-mouse Sca-1 to label murine megakaryocytes and HSPCs, respectively. The immunostained tissues were secondarily stained with Alexa Fluor 488 anti-rabbit -or- Alexa Fluor 405 anti-rat, to label CD41^+^ and Sca-1^+^ cells, respectively. Finally, the prepared samples were imaged via confocal microscopy (Invitrogen EVOS M7000 and Zeiss LSM 880).

## Supporting Information

Supporting Information is available from the Wiley Online Library or from the author.

## Author contributions

Each author’s contribution(s) to the paper are listed below:

Conceptualization: ETP

Methodology: SD, WT

Investigation: SD, WT, ETP

Visualization: SD, ETP

Supervision: ETP

Writing—original draft: SD

Writing—review & editing: SD, WT, ETP

## Acknowledgements

We would like to thank Christina Stinger from the University of Delaware Office of Laboratory Medicine for assistance with the animal experiments. We would also like to thank the members of the Nemours Histology Core Facility and Charles Riley from the Delaware Biotechnology Institute for their assistance with the tissue sectioning and immunostaining. Finally, we would like to thank Shannon Modla and Jean Ross and other members from the Bioimaging Center at the Delaware Biotechnology Institute for their assistance with transmission electron microscopy. This work was supported in part by a grant by CSL Behring through the Science Center (Philadelphia, PA, USA; https://sciencecenter.org) and the US National Science Foundation, NSF (CBET-1804741). Microscopy equipment (Carl Zeiss LSM880 confocal microscope) was acquired with a shared instrumentation grant (S10 OD016361) and access was supported by the NIH-NIGMS (P20 GM103446), the US NSF (IIA-1301765) and the State of Delaware. Use of the University of Delaware Animal Laboratory and Histology Core Facility were supported by the DCMR COBRE program, with a grant from the National Institute of General Medical Sciences and from the National Institutes of Health – NIH-NIGMS COBRE (P20 GM139760).

## Ethics Statement

Mice were handled humanely under IACUC approved Animal Use Protocol #1290.

