## Supplemental Material for "Native and engineered human megakaryocytic extracellular vesicles for targeted non-viral cargo delivery to blood stem cells"

**Address**: Ammon-Pinizzotto Biopharmaceutical Innovation Building
Delaware Biotechnology Institute
590 Avenue 1743
Newark, DE 19713, USA

**Key Words:** hematopoietic stem & progenitor cells (HSPCs), targeted delivery, gene therapy, tropism, megakaryocytes, platelets, biodistribution

| Experiment | Dose | 4-hours | 24-hours | 48-hours | 96-hours |
| --- | --- | --- | --- | --- | --- |
| Saline (PBS)-treated mice (Female NSG \| 7-8-weeks old) | | | | | |
| Blood Platelet Analysis | 150-µL | 3 | 4 | 4 | 4 |
| Biodistribution *(Includes mice from platelet analysis)* | 150-µL | 2 | 3 | 3 | 3 |
| huMkEV-treated (PKH26-stained) mice (Female NSG \| 7-8-weeks old) | | | | | |
| Blood Platelet Analysis | $6{\times}10^6$ EVs | 6 | 9 | 5 | 6 |
| Biodistribution *(Includes mice from platelet analysis)* | $6{\times}10^6$ EVs | 5 | 9 | 5 | 5 |
| Total treated mice per timepoint (huMkEV + PBS): | | 9 | 13 | 9 | 10 |

**Supplemental Table S1.** Mice counts per condition. Count of 7-8-week female NSG mice for assessing *in vivo* huMkEV-induced megakaryopoiesis and EV biodistribution at 4-, 24-, 48-, and 96-hours following intravenous administration.

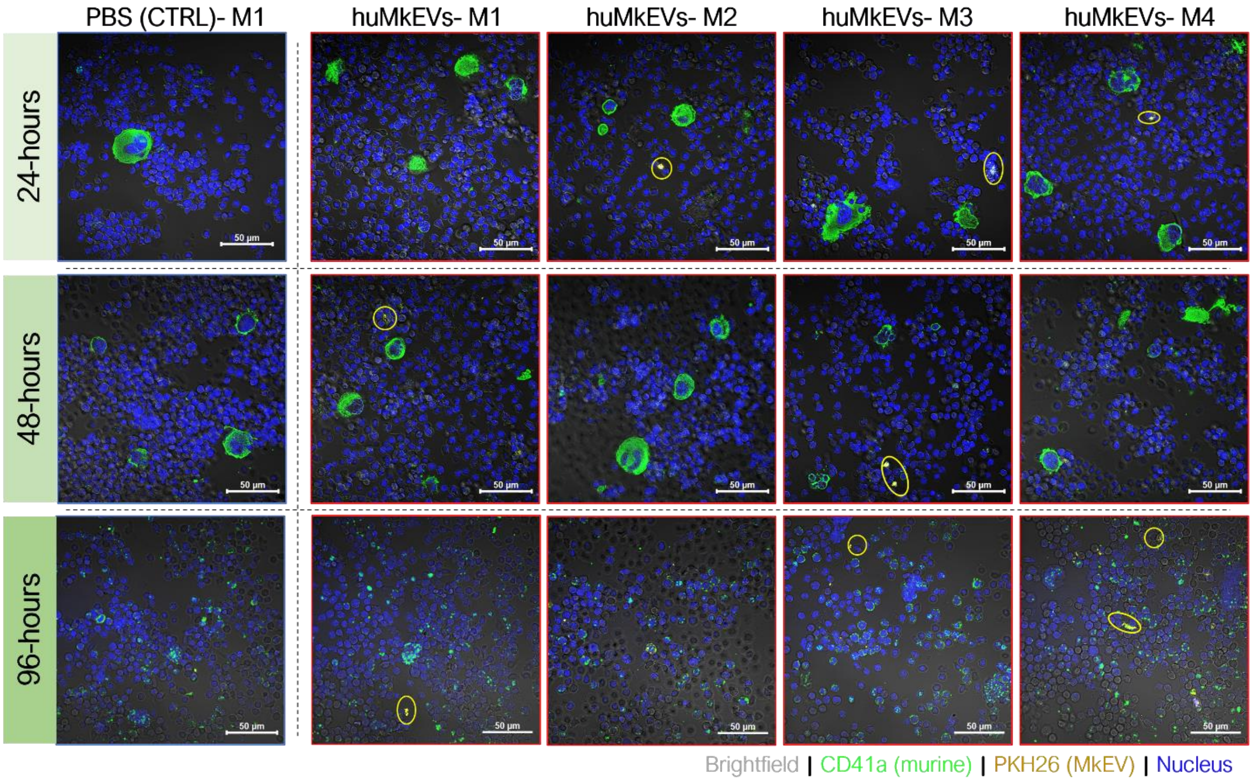


**Supplemental Figure S2.** Development of mature megakaryocytes in murine bone marrow following huMkEV treatment. Select bone marrow flushes from PBS-treated (left column) and huMkEV-treated (right 4 columns) mice from 24-hours (top row), 48-hours (middle row), and 96-hours (bottom row) were immunostained for murine CD41a (green) to determine the degree of differentiation of the murine HSPCs to the megakaryocytic phenotype. Presence of huMkMPs indicated in yellow (PKH26) and nuclei are shown in blue (DAPI). Scale bars: 50-µm.

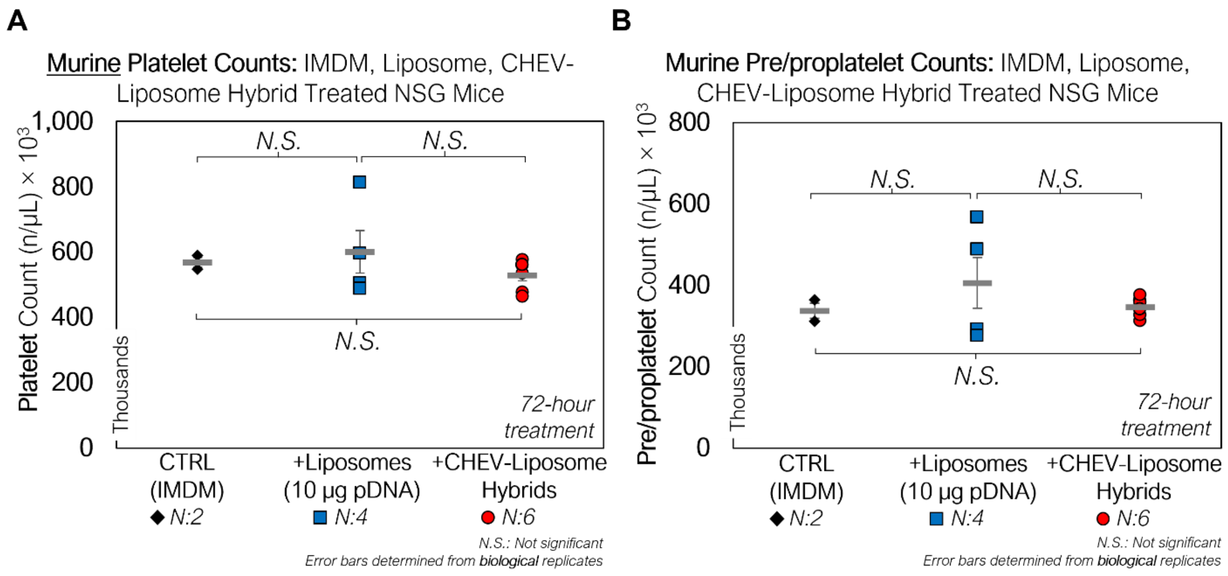


**Supplemental Figure S3.** Impact of liposome, CHEV-liposome hybrid treatment on platelet counts. A) Platelet and B) pre/proplatelet counts in NSG peripheral blood 72-hours following treatment with IMDM (media control), pDNA-loaded liposomes, or CHRF EV-liposome hybrids. *N.S.: not significant, Student's T-test*

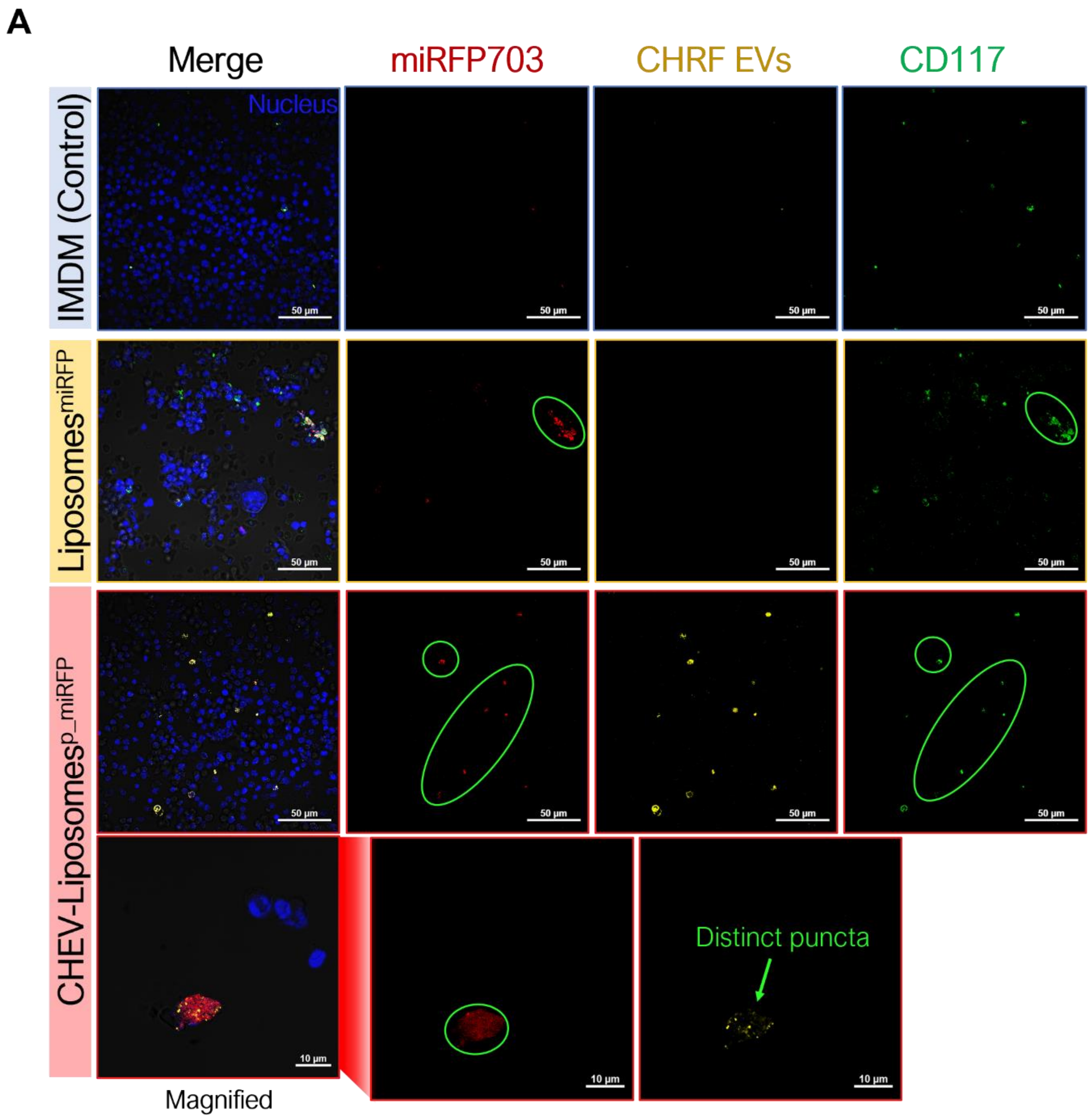


**Supplemental Figure S4.** Flushed marrow (femur) cells from mice treated with pDNA –loaded CHEV-liposome hybrids exhibit expression of miRFP703 from delivered plasmid. Flushed RBC-depleted marrow cells from mice treated with either pDNA-loaded liposomes (pLifeAct-miRFP703) or pDNA-loaded CHEV-liposome hybrids and a media (IMDM)-treated control were immunostained for CD117+ HSPCs (green) and screened for miRFP703 expression (red), and CHEVs (yellow). Colocalization of murine HSPCs and miRFP703 indicated with light green circles and nuclei are shown in blue.

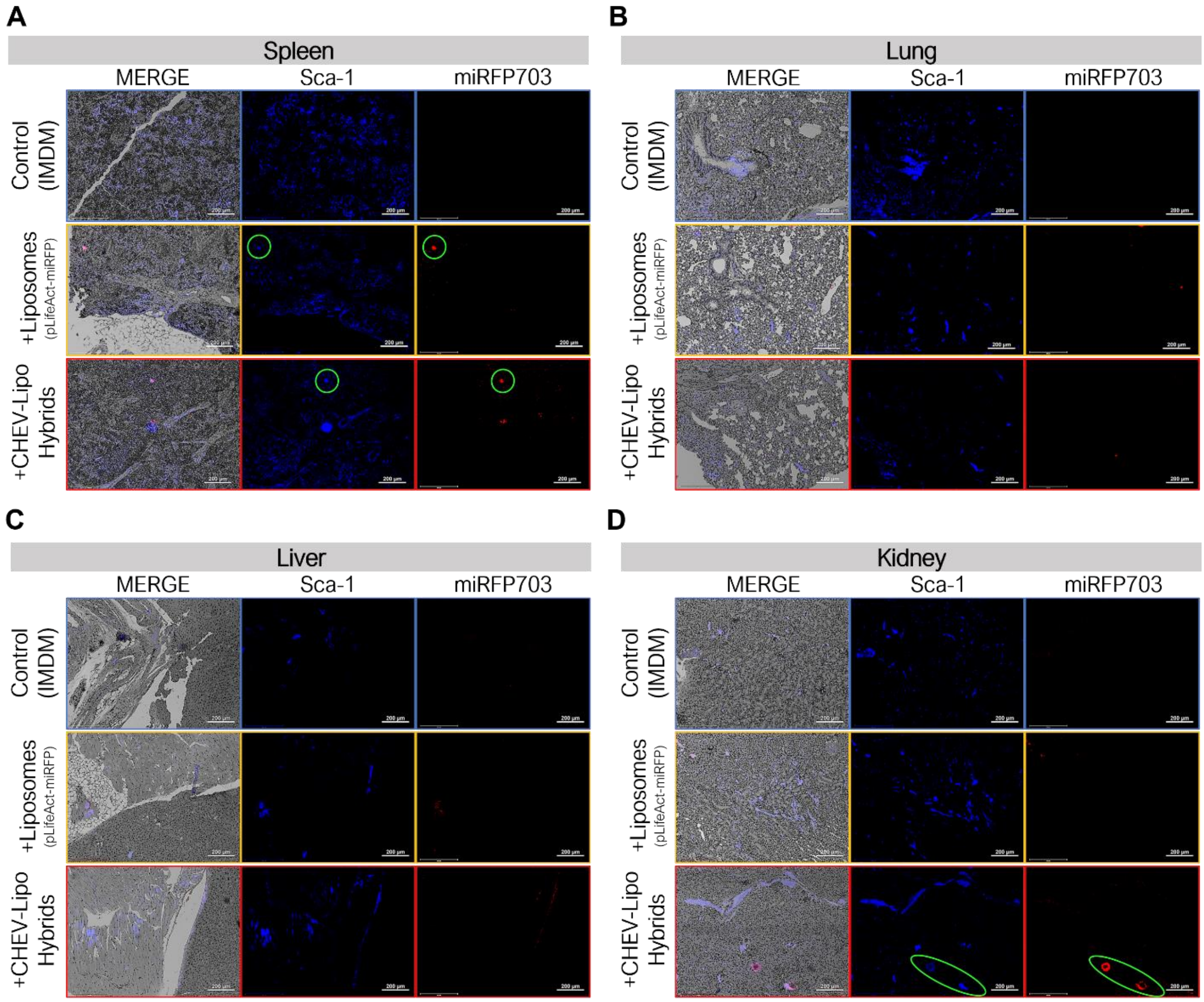


**Supplemental Figure S5.** Histological analysis of various Sca-1 immunostained tissues excised from pDNA-loaded liposome and CHEV-liposome hybrid-treated mice. Lungs, spleen, kidneys, and liver from 72-hour-treated mice were fixed (10% neutral-buffered formalin), sectioned, immunostained and assessed for structure (gray- DIC) and presence of Sca-1+ cells (blue), miRFP703 expression (red) and PKH26 arising from administered CHEV hybrids. Colocalization of murine HSPCs and miRFP703 indicated with light green circles. Scale bars: 200-µm.

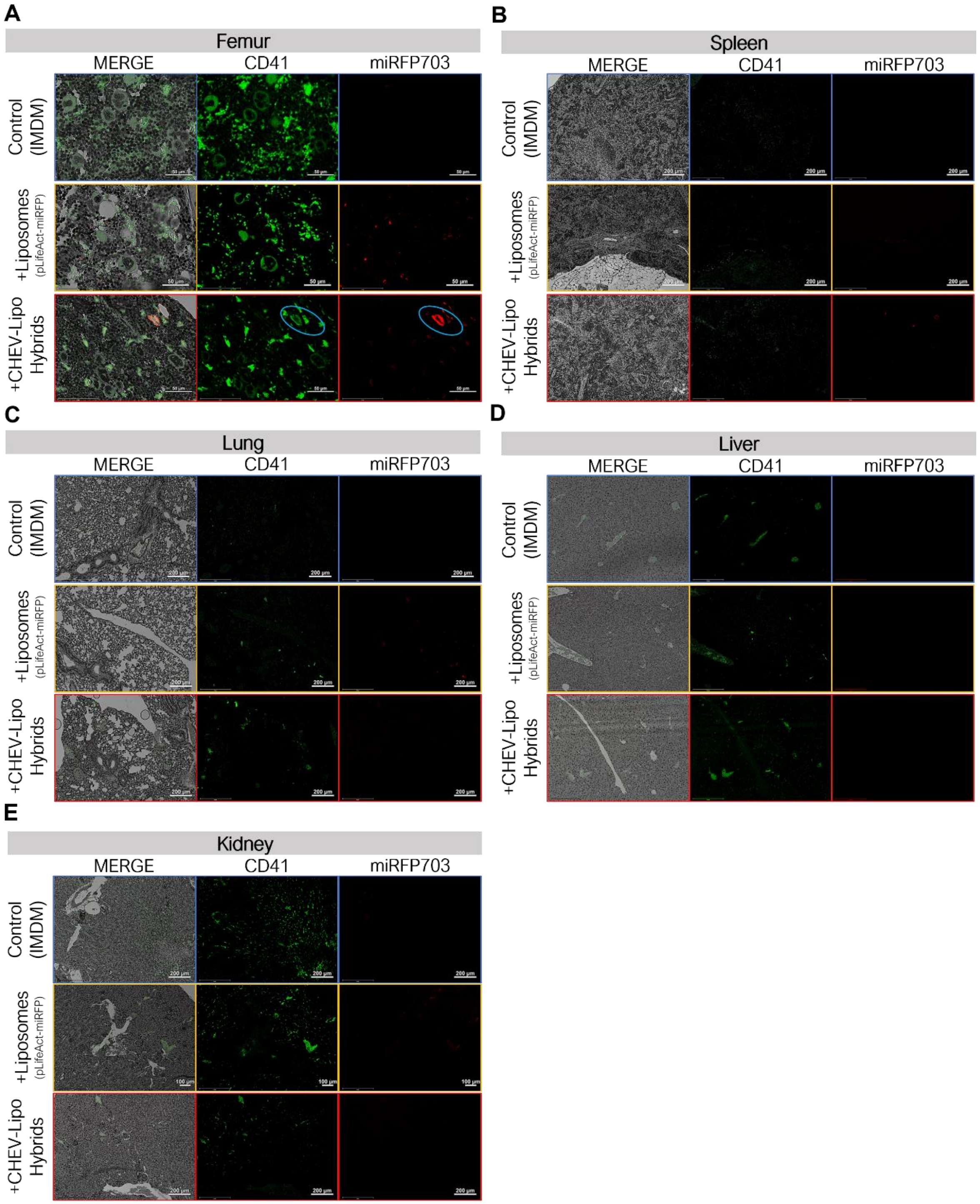


**Supplemental Figure S6.** Histological analysis of various CD41 immunostained tissues excised from pDNA-loaded liposome and CHEV-liposome hybrid-treated mice. Lungs, spleen, kidneys, and liver from 72-hour-treated mice were fixed (10% neutral-buffered formalin), sectioned, immunostained and assessed for structure (gray- DIC) and presence of CD41+ cells (green), miRFP703 expression (red) and PKH26 arising from administered CHEV hybrids. Colocalization of murine CD41+ megakaryocytes and miRFP703 indicated with light blue circles. Scale bars: 200-µm.
